# Unraveling reproducible dynamic states of individual brain functional parcellation

**DOI:** 10.1101/2020.03.02.972760

**Authors:** Amal Boukhdhir, Yu Zhang, Max Mignotte, Pierre Bellec

**Affiliations:** Centre de recherche de l’institut universitaire de gériatrie de Montréal, Montréal, Québec, Canada; Département d’informatique et de recherche opérationnelle, Université de Montréal, Montréal, Québec, Canada; Département de psychologie, Université de Montréal, Montréal, Québec, Canada

## Abstract

Data-driven parcellations are widely used for exploring the functional organization of the brain, and also for reducing the high dimensionality of fMRI data. Despite the flurry of methods proposed in the literature, functional brain parcellations are not highly reproducible at the level of individual subjects, even with very long acquisitions. Some brain areas are also more difficult to parcellate than others, with association heteromodal cortices being the most challenging. An important limitation of classical parcellations is that they are static, i.e. they neglect dynamic reconfigurations of brain networks. In this paper, we proposed a new method to identify dynamic states of parcellations, which we hypothesized would improve reproducibility over static parcellation approaches. For a series of seed voxels in the brain, we applied a cluster analysis to regroup short (3 minutes) time windows into “states” with highly similar seed parcels. We splitted individual time series of the Midnight scan club sample into two independent sets of 2.5 hours (test and retest). We found that average within-state parcellations, called stability maps, were highly reproducible (over .9 test-retest spatial correlation in many instances) and subject specific (fingerprinting accuracy over 70% on average) between test and retest. Consistent with our hypothesis, seeds in heteromodal cortices (posterior and anterior cingulate) showed a richer repertoire of states than unimodal (visual) cortex. Taken together, our results indicate that static functional parcellations are incorrectly averaging well-defined and distinct dynamic states of brain parcellations. This work calls to revisit previous methods based on static parcellations, which includes the majority of published network analyses of fMRI data. Our method may, thus, impact how researchers model the rich interactions between brain networks in health and disease.

## 1. Introduction

Brain parcellation is a tool for understanding the functional organization of the human cerebral cortex, and also to reduce the dimensionality of fMRI data. Parcellations are notably heavily used to characterize brain network properties. A brain parcellation was defined as the entire subdivision of the brain into clusters (or spatially distributed parcels/regions). A good parcellation should typically satisfy two conflicting objectives. The first objective is to be reproducible enough to allow for replication and comparison across studies. The second objective is to be flexible enough to accurately represent the organization of an individual brain. Simultaneously achieving these two objectives is challenging, in part due to inter-subject variability (Braga & Buckner, 2017; Nastase et al., n.d.; Sripada et al., 2019), which is also associated with measures of cognitive performance (Betzel et al., n.d.). In addition, it has also become apparent that brain functional connectivity substantially reorganizes dynamically (Yaesoubi et al., 2018) according to different cognitive states (Salehi, Greene, et al., 2018). Our main objective in this work was to develop a new method to capture reproducible dynamic states of parcellations at the individual level. These dynamic states represent the spatial brain reconfigurations over time in the resting state condition.

Even though there has been remarkable progress in the field of brain parcellation, there are still concerns about the reproducibility of individual level parcellations. For instance, Gordon and colleagues confirmed that subject-to-group similarity was relatively low (Gordon, Laumann, Adeyemo, et al., 2017). This was measured by the average parcel connectivity similarity to group connectivity score, which reached a plateau with an average of 0.7 for long scan duration (∼ 800 timepoints) (Gordon, Laumann, Gilmore, et al., 2017). Researchers reported that the lateral prefrontal cortex and the lateral temporal occipital cortex were among the most variable parcels across subjects (Marek & Dosenbach, 2018). We hypothesize that the main limitation of these approaches was the use of a fixed parcellation over time, which neglected the dynamic aspect of heteromodal brain networks (Salehi, Greene, et al., 2018). In other words, different brain reconfigurations for each parcel were forced to be averaged in one brain parcel while suppressing region boundaries reconfigurations over time. In the literature, Yeo and colleagues reported results in line with our hypothesis, showing that brain networks were continually aggregating and segregating over time (Yeo et al., 2014). This suggests the importance of considering the dynamic aspects of brain parcellation, rather than considering it as a fixed system.

A proliferation of approaches also exist in the literature to study the dynamics of functional connectivity. These studies confirmed the spatio-temporal reconfiguration of the brain networks (Allen et al., 2014; Vince D. Calhoun et al., 2014; Donnelly-Kehoe et al., 2019; Hutchison et al., 2013) and associated it to dynamics of cognitive processing or mental states dictated by tasks (Bassett et al., 2011; Braun et al., 2016; Gonzalez-Castillo et al., 2015; J. Liu et al., 2018; Reinen et al., 2018). The co-activation patterns (Chen et al., 2015; X. Liu & Duyn, 2013), spatial independent component analysis (Smith et al., 2012) were among the most widely applied techniques to consider brain dynamics (see (Chen et al., 2017) for a review). These dynamic analyses of the brain, using for instance sliding window correlation, have demonstrated better results compared to stationary approaches in the detection of neurological disease (Iraji et al., 2018; Sakoğlu et al., 2010). Other findings confirmed the interaction between brain networks for different task states (Braga & Buckner, 2017; Casorso et al., 2019; Yeo et al., 2014). For instance, Braga and Buckner discovered that the default mode network could be reliably subdivided into parallel networks within the same individual (Braga & Buckner, 2017). Chen and colleagues modeled these states switching processes of resting state brain activities using a hidden markov model (Chen et al., 2015). Therefore, neuroscientists mentioned there is a need to have neuroimaging tools to identify how brain parcels reconfigure spatially in the case of highly cognitive regions over time and to evaluate the variability of brain parcels across time and across individuals (J. Liu et al., 2018). Even though dynamic functional connectivity is well studied, to the best of our knowledge, the only parcellation approach that considered dynamic changes of parcels was suggested by Salehi and colleagues. These authors demonstrated that the brain functional parcellations are not spatially fixed, but reconfigure with task conditions (Salehi, Greene, et al., 2018; Salehi, Karbasi, et al., 2018). These reconfigurations were used to reliably predict different task conditions. Still, this approach only suggested a brain parcellation per task condition, and it neglected brain parcel reconfigurations across short time durations, within each task. Dynamic brain parcellation, thus represents a promising area of research to further investigate brain dynamics.

In this paper, we build upon the findings of Salehi and colleagues (Salehi, Greene, et al., 2018), and we propose a novel approach to extract different dynamic states of functional parcellations at the individual level. We define a dynamic state of parcellation as the spatial reconfiguration of a given brain network that occurs for short time durations in the resting state condition. We hypothesize the existence of homogeneous modes of spatial reconfigurations, or dynamic states of parcellations, at the level of these short time windows and we propose a dynamic cluster analysis for their identification. Our approach is based on aggregating sliding-window parcellations for a given region to obtain stability maps of the different dynamic states of parcellations. We generate these dynamic states for the ten subjects of the Midnight scan club (MSC) resting-state dataset and we aim to study similarities and variations within-state (across replication sets), across states (within-subject), and across subjects. We also aim to evaluate the reliability of the generated states maps in a “fingerprinting” experiment, i.e. matching state maps generated from the same subjects within a group.

## 2. Materials and methods

### 2.1 Dataset and preprocessing

The resting-state MSC dataset includes ten healthy subjects (female=5, male=5, their age ranges between 24-34 years old (Gordon, Laumann, Gilmore, et al., 2017). Informed consent was obtained from all participants. The study was approved by the Washington university school of medicine human studies committee and institutional review board (Gordon, Laumann, Gilmore, et al., 2017). Each subject underwent a total of five hours of resting state functional MRI data, with a series of 30 minutes contiguous acquisitions, beginning at midnight for ten consecutive days. In each session, subjects visually fixated on a white crosshair presented against a black background. All functional imaging was performed using a gradient-echo EPI sequence (TR = 2.2s, TE = 27 ms, flip angle = 90°, voxel size = 4 mm × 4 mm × 4 mm, 36 slices) on a Siemens TRIO 3T MRI scanner. An EyeLink 1000 eye-tracking system allowed continuous monitoring of the eyes of the subjects in order to check for periods of prolonged eye closure, potentially indicating sleep. Only one subject (MSC08) demonstrated prolonged eye closures. For details about the data acquisition parameters see (Gordon, Laumann, Gilmore, et al., 2017). The MSC dataset was preprocessed and analyzed using the <u>NeuroImaging Analysis Kit</u>^1^ executed within a Ubuntu 16.0.4 Singularity container, running GNU Octave version 4.2.1, and the MINC toolkit version 1.9.15. The first five volumes of each run were suppressed to allow the magnetisation and reach equilibrium. Time series were normalized to the zero mean and unit variance. Each fMRI session was corrected for inter-slice difference in acquisition time and the parameters of a rigid-body motion was estimated for each time frame. The “scrubbing” method of (Power et al., 2012), was used to remove the volumes with excessive motion (frame displacement greater than 0.5). No session was excluded due to excessive motion. Each session had at least 420 volumes after scrubbing, across all subjects, and with a maximum of 810 volumes available. Also, the nuisance parameters were regressed out from the time series at each voxel i.e., slow time drifts, average signals in conservative masks of the white matter and the lateral ventricles, as well as the first principal components of the six rigid-body motion parameters and their squares. The fMRI volumes were spatially smoothed with a 6 mm isotropic Gaussian blurring kernel. A more detailed description of the preprocessing pipeline can be found on the NIAK website ^2^.

### 2.2 Individual dynamic states of parcellation

We developed an algorithm which identifies dynamic states of brain parcellation at the individual level called Dynamic Parcel Aggregation with Clustering (Dypac). The algorithm is composed of four steps as illustrated in Fig. 1. In the first step (Fig. 1A), we select a series of sliding time windows from individual fMRI time series (*W=100* time points, with *O=10* time points of overlap, starting from the first time point), and we generate parcellations for the whole cerebral cortex using a k-Means clustering algorithm (Pedregosa et al., 2011). We select fMRI time windows from several runs, such that some time series may combine signals from separate runs. The motivation behind this parcellation step is the identification of brain regions with similar temporal activity for a given time window. We choose the k-Means for its linear complexity and simplicity to run; i.e. no need for many parameters to tune (David Arthur Stanford University, Stanford, CA & Sergei Vassilvitskii Stanford University, Stanford, CA, n.d.; MacQueen, 1967). Our algorithm is parallelized based on the multiprocessing library to run computations on multiple cores^3^. We also used k-Means using scikit-learn implementation^4^ (Pedregosa et al., 2011) to generate parcellations and used the k-Means++ method to choose initial cluster centers in a strategic way in order to speed-up convergence. Although the k-Means algorithm has the drawback of falling into local minima, the k-Means++ initialization helps overcome this limitation with a better exploration of the parcellation solution search space. We also replicate the k-Means parcellations (repetition = 5) for each window with different initializations of the random number generator. This helps identify consistent solutions across different local minima. All the replicated solutions with different seeds are pooled with the set of k-Means parcellations. That is, we simply used all the K-Means based parcellations from different sliding windows as an input for the similarity matrix of the Hierarchical clustering. The total number of K-Means parcellations used in the similarity matrix equals the number of sliding windows multiplied by the number of replications. This result is multiplied by the number of sessions (e.g. if the number of replications=5, number of sliding windows=10, number of sessions=2 then the total number of K-Means parcellations = 100).

**Figure 1.**
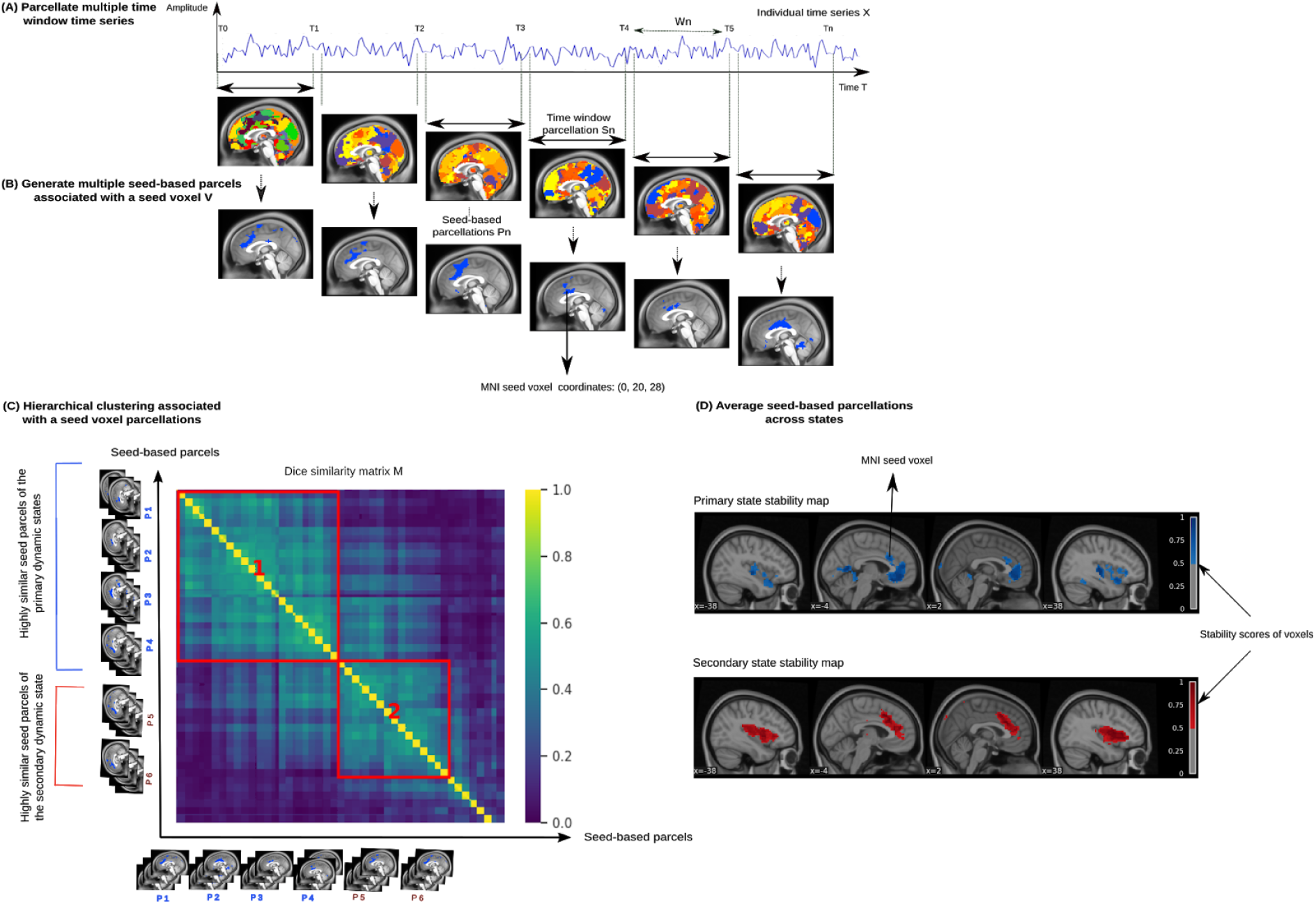
Subject-specific dynamic parcellation approach (Dypac); (A) we generate multiple short time window parcellations for the whole cerebral cortex per subject. (B) Then, we identify the parcel associated with a particular seed-voxel. (C) We calculate the pairwise similarity of these seed-based parcellations based on the Dice similarity score. We apply a Hierarchical clustering on this similarity matrix to group these parcellations into a set of clusters (or dynamic states of parcellations) according to a threshold imposed on the Dice similarity matrix, as well as the number of sliding windows included in the state. (D) For each cluster and each seed, we average all of its seed-based parcellations to obtain the final dynamic state stability maps.

In the second step (Fig. 1B), we identify for a particular seed voxel which parcel this voxel is associated with. We thus obtain a binary representation called seed-based parcellation. This step may contribute to the success of our approach, since it allows us to simplify the complexity of the full brain parcellation problem by focusing on the functional activity of one region of interest.

In the third step (Fig. 1C), we calculate the pairwise similarity of all seed-based parcellations generated from different sliding windows. A seed-based parcellation was defined as the subdivision of the entire subnetwork, associated with a given seed, into spatial clusters (i.e., spatially distributed regions/parcels). This similarity is measured with the so-called Dice similarity score. Then, we apply the Hierarchical clustering (here, with the average linkage method) on the Dice-similarity matrix to group seed-based parcellations into dynamic clusters or ‘states’, according to an empirical similarity threshold. This threshold constrains the clustering of seed-based parcellations by requiring a minimum Dice similarity (here, 0.3) between these parcellations in the same dynamic state, which will in turn infer the number of states in a data-driven way. Given the previously mentioned settings, we hypothesize that some dynamic states of parcellations only appeared in few sliding windows, i.e. inferior to 10% of the total number of available seed-based parcellations. These states might be associated with spurious or non-reproducible spatial brain reconfigurations. Then, we apply another threshold which filtered the identified states based on their number of seed-based parcellations in order to keep only those with more than 10% of the total number of seed-based parcellations. This second threshold will remove noisy states in order to keep reproducible patterns over time. There should be a trade-off between the similarity threshold imposed on the Dice similarity matrix and the threshold that constrains the number of seed-based parcellations in a given state. In other words, a higher Dice score threshold allows to obtain smaller states (i.e., the higher the Dice score, the lower the number of seed-based parcellations of a given state) and thus, it requires a smaller number of seed-based parcellations threshold. This allows us to avoid missing the most interesting states.

In the last step (Fig. 1D), we averaged all seed-based parcellations for a given cluster to get its state stability map. This provides a probability of each voxel to be assigned to a given state as a measure of the stability of voxels with respect to their membership in a particular area. Stability maps represent the spatial signature of each dynamic state of brain parcellations, and the final outcome of the algorithm.

Our method generates dynamic states of parcellations as functionally distributed subnetworks across the brain or local subnetworks surrounding the seed. We also suggest a simple conversion to split subnetworks into multiple spatially contiguous/connected regions instead of spatially distributed parcels. This can be useful in the context of graph theory by considering contiguous regions as nodes in the graph. To this end, we apply a constraint that separates out connected components and assigns to each region a unique state label, using the Nilearn function implementation^5^. We set the minimum region size in volume required to keep after extraction to 50 voxels. This removes small or spurious regions.

### 2.3 Choice of the studied subnetworks and their seed

To generate seed-based parcellations, we studied seed voxels from three regions of the MIST parcellation. We picked MNI coordinates (0, -76, 10), (0, 20, 28) and (3, -43, 37) as the respective medoids of the ROIs 90, 6 and 42 corresponding to the posterior medial visual subnetwork (PM-VIS), the dorsal anterior cingulate cortex (dACC) and the posterior cingulate cortex (PCC) subnetworks in the MIST atlas (Urchs et al., 2017). The choice of seeds was driven by the properties of the networks in the literature. We first chose a seed from an area with the least functional variability (Gordon et al., 2016): PM-VIS, a core visual area (Gordon, Laumann, Adeyemo, et al., 2017). In the case of the dACC, this region played a prominent role within the salience network, which is involved in many functions including response selection, conflict resolution and cognitive control and it is among the most highly dissimilar networks across subjects (Gordon et al., 2016; V. Menon, 2015; Vinod Menon & Uddin, 2010). Finally, the PCC is considered as a hub node in the default mode subnetwork. Previous findings reported it as a highly heterogeneous network and suggested it may play a direct role in regulating the focus of attention, memory retrieval, conscious awareness and future planning. Also, functional interaction between the nodes of the salience seed and those of the default mode, including the PCC, during moral reasoning is reported in previous studies (Chiong et al., 2013; Jilka et al., 2014).

### 2.4 Spatial reproducibility analysis

To evaluate the quality of the dynamic states of parcellations, we conducted two quantitative analyses. First, we compared the performances of the Dypac algorithm with a static parcellation algorithm; i.e. the k-Means algorithm. This allowed us to compare the goodness of our dynamic states of parcellations with an existing static state of the art parcellations. Second, we conducted a quantitative consistency analysis at the within-subject level. This allowed us to identify, for a given subject, similarities and variations in the spatial reconfigurations across states and seeds. We conducted a reproducibility analysis for the two analyses.

In both analyses, we half-split the Midnight scan club dataset into two equally sized sets of five independent sessions (of a total of 2.5 hours each) per subject. Each half (about 2.5 hours per subject) was used to replicate several seed-based parcellations that we called replication sets. Then, we generated dynamic states of parcellations based on our proposed approach (See Fig. 1). We sorted the states by decreasing dwell time for the first replication set (i.e. the cumulative durations of all sliding windows associated with a given state, relative to the total duration of the scan). Accordingly, we labeled the states of the first replication set into primary state, secondary state, tertiary state, etc. Prior to comparing our state stability maps, we matched the first set maps to maps from the second set using the Hungarian method (Kuhn, 2005) which used the Pearson correlation for the spatial matching between state stability maps. The Hungarian method was applied to the within-subject and between-subjects analysis. A high correlation reflected a strong linear relationship between states maps and is indicative of consistent spatial regions from the two sets of independent data. We replicated these consistency analyses across all states and all subjects of the Midnight scan club dataset. We run both our Dypac algorithm and the K-Means algorithm 15 times per set with different random seeds due to the stochastic aspect of the k-Means algorithm. This allowed us to verify the sensitivity of both algorithms to local minima.

### 2.5 Fingerprinting experiment

We finally evaluated the individual specificity of our dynamic states of parcellations by attempting to match dynamic states maps generated from data acquired on the same subjects, when these maps are mixed with maps generated from other subjects. To this end, we cross-correlated a given state stability map with all state stability maps from all subjects.

The state stability maps were generated from the split half sets of the Midnight scan club dataset and all these maps were pooled altogether in the fingerprinting. For a given map, we looked for the map that matched the closest map from the pool of all maps. Each seed subnetwork was analyzed separately. A fingerprinting was successful when maximal correlation was observed between a pair of two state stability maps originating from the same subject, otherwise it was considered as a failure. We repeated this experiment for all state maps across all subjects and seeds. We denoted this procedure by the deterministic fingerprinting. We computed the accuracy score as the number of correct states matching over the total number of matched maps. A high accuracy score revealed that state stability maps were reliable to differentiate subjects based on their specific spatial brain reconfigurations. Inversely, a low accuracy score was associated with state stability maps which were either very similar across subjects or unreliable within subjects.

To correct for the different number of identified states for each subject, we run a fingerprinting by chance experiment. To do that, we selected a state map, arbitrarily for each subject. Second, we selected another second state map arbitrarily from the pool of state stability maps of all subjects. If these two maps belonged to the same subject, then the fingerprinting was successful, otherwise it was considered a failure. We repeated this process 1000 times. We computed the accuracy of the fingerprinting by chance and compared it to the deterministic fingerprinting.

### 2.6 States dwell time reproducibility analysis

We aimed to get a better understanding of the dwell time reproducibility over time of the dynamic states; i.e., the proportion of the total number of sliding windows that were associated with a given state. To this end, we performed a spatial matching of states between two sets of independent sessions in terms of the Pearson correlation and we reported their associated dwell times in Fig. 12. This matching was based on the Hungarian method. Therefore, only the dwell times associated with spatially reproducible states were included.

### 2.7 Data records

Scripts used in this study are available on Github^6^. The generation of state stability maps can be executed online via a Jupyter notebook via the binder platform. We have also made available online all the state stability maps for the ten subjects of the MSC dataset on the neurovault website^7^.

## 3. Results

### 3.1 Temporal cluster analysis reveals “dynamic parcellation states” with highly homogeneous parcellations within a state, and highly dissimilar parcellations across states

We first aimed to assess whether homogeneous parcellations can be extracted from short time windows of about 3 minutes duration. We replicated seed-based parcellations on 220 sliding-windows for this purpose. These time windows were extracted from a pool of time samples, generated by randomly concatenating five sessions of imaging data for the Midnight scan club sample, resulting in a total of 2.5 hours of fMRI signals per subject.

For a given seed voxel in the brain, we observed pairs of seed-based parcellations with high homogeneity across different time windows: Dice coefficients between pairs of seed-based parcellations were larger than 0.8, or 0.9 for some seeds and subjects, see Fig. 2. We reported the Dice coefficients distributions for the identified states across the studied subnetworks (see Supplementary Material 3). By contrast, many pairs of seed-based parcellations associated with different sliding windows had very low Dice scores, close to zero, despite being associated with the same seed. For example, in the similarity matrix of subject MSC02, bright colors were associated with some highly homogeneous seed-based parcellations across the diagonal, while the remaining pairs of seed-based parcellations were associated with low Dice scores (blue color). This observation motivated us to develop a “dynamic cluster analysis”, grouping seed-based parcellations on sliding windows into a number of homogeneous ‘dynamic states of parcellations’, for a given seed voxel. This approach allowed us to disentangle different dynamic states of parcellations based on the variability of their spatial distribution over time, specifically for a given brain subnetwork. Moreover, our findings suggested the existence of different temporal dynamics for most of the different states either associated with the same seed or different seeds (See Supplementary Materials 6).

**Figure 2.**
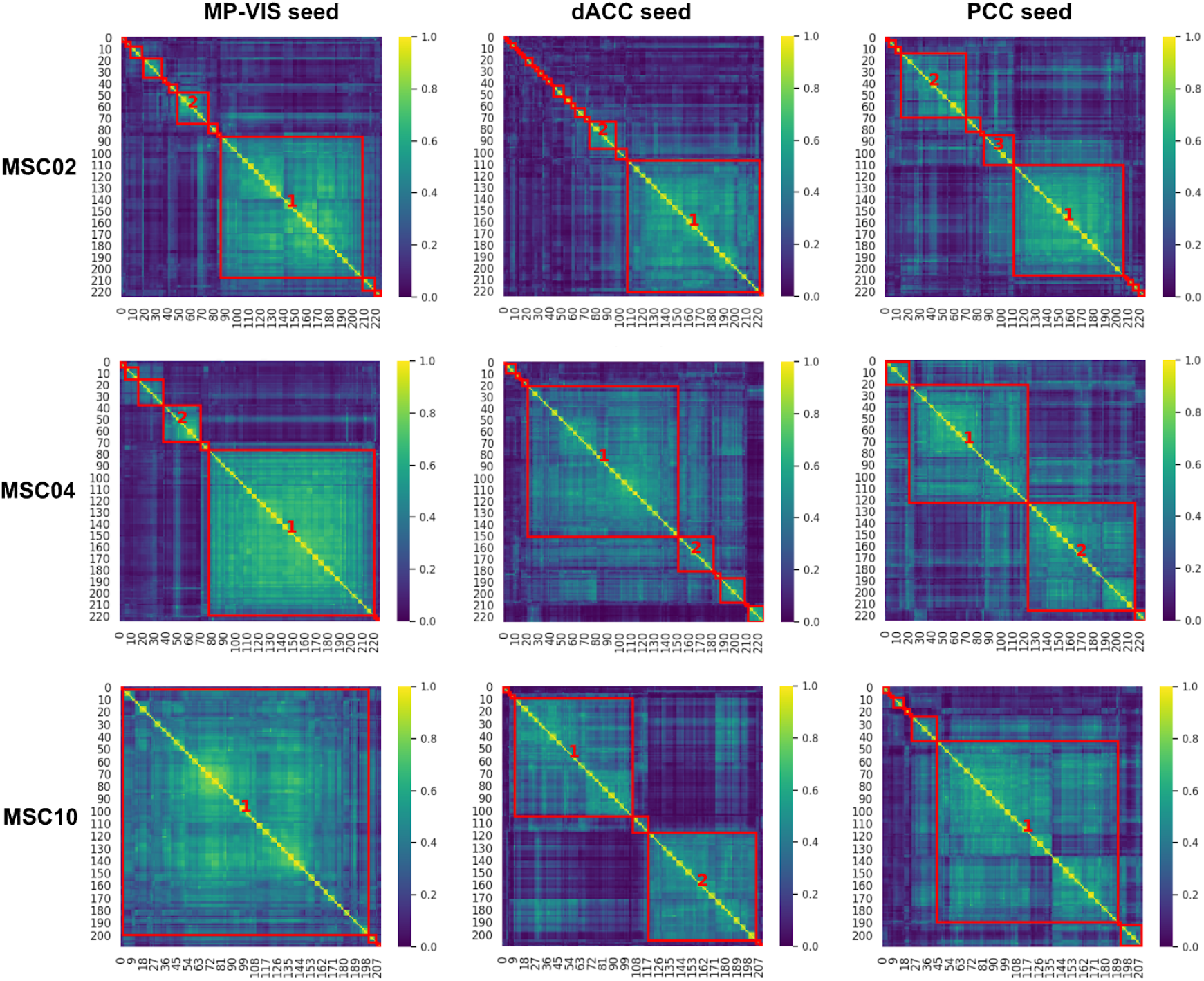
Dice similarity matrices of seed-based brain parcellations showed groups of highly homogeneous seed-based parcellations and other groups of dissimilar seed-based parcellations. Each element in the similarity matrix represented the Dice score between a pair of seed-based parcellations. This matrix was calculated, separately for each subject and each seed voxel. Three subjects of the MSC dataset and three seeds were investigated, i.e. the Posterior Medial Visual subnetwork (PM-VIS), the dorsal Anterior Cingulate (dACC) and the Posterior Cingulate Cortex (PCC).

Each dynamic state was characterized by its dwell time relative to the total scan duration, i.e. the proportion of the total number of sliding windows that were associated with a given state. We applied two criteria to decide on the number of states for a seed voxel: 1) seed-based parcellations within a state had to exhibit a minimal average level of Dice similarity; i.e. Dice > 0.3, and 2) the dwell time of a given dynamic state needed to be substantial, i.e. larger than 10%. For example, using these two criteria for the PCC seed and subject MSC02, three separate dynamic states of parcellations were identified, and together these dynamic states of parcellations added to about 75% dwell time of all available sliding-windows, see Fig. 2.

For a better understanding of the dwell time distribution across dynamic states, we showed its distribution across the ten subjects of the Midnight scan club dataset and the Dypac algorithm. The results showed the existence of a dominant state for the three studied subnetworks. For instance, the primary states of the PM-VIS had a median dwell time of 73% over only 11% median dwell time in the case of its secondary states. Similarly, the primary states of the PCC had a median dwell time of 63% over only 20% median dwell time in the case of its secondary states. Moreover, the dACC and the PCC subnetworks were multistate with up to five states in the case of the dACC seed and up to four states in the case of the PCC seed. Less states were observed in the case of PM-VIS with very low dwell time, i.e. ∼10%. Therefore, the PM-VIS was monostate for most subjects even though some subjects had multistate maps with a dominant primary state (See Fig. 3).

**Figure 3.**
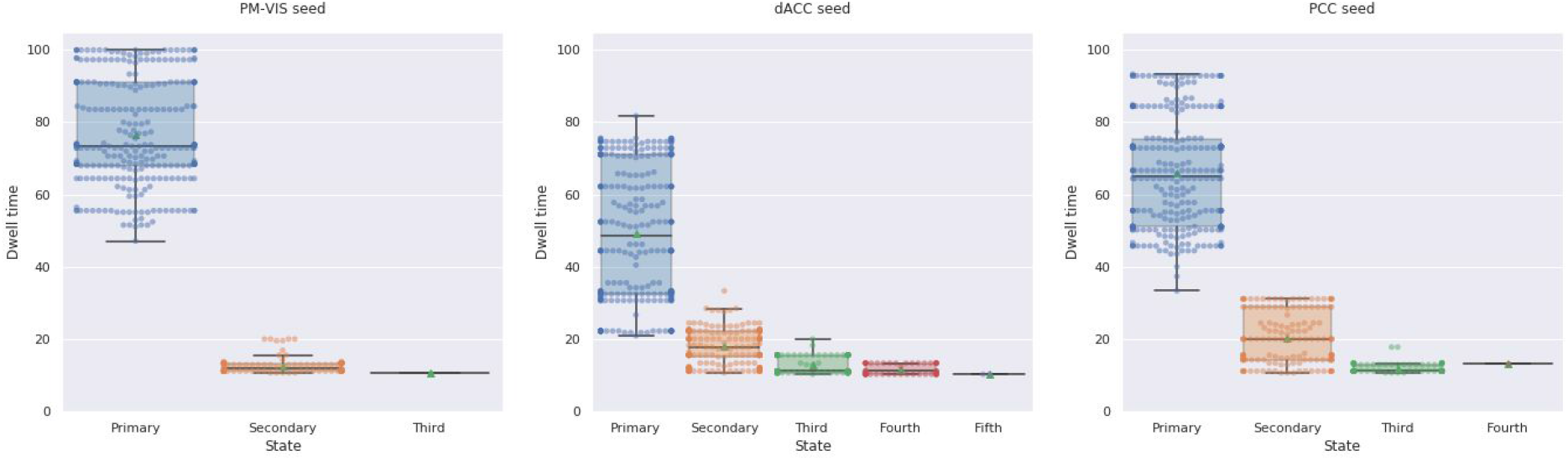
Subnetworks were multistate with a dominant primary state. We included state dwell times for both sets of independent data. Three seed subnetworks were investigated including the PM-VIS, the dACC and the PCC subnetworks across ten subjects of the Midnight scan club dataset. The number of replications of seed-based parcellations per sliding window = 5. We reported dwell times from 30 replications of the Dypac algorithm.

Taken together, these results showed that resting-state brain parcellations fall into a number of highly homogeneous and distinct states over time, at an individual level. We also investigated the impact of different parameters of our Dypac algorithm on the states dwell time including the window length (Supplementary Material 4.3), the cluster size threshold (Supplementary Material 4.5) and the smoothing kernel size (Supplementary Material 4.6). Our findings were consistent across different parameters.

### 3.2 Dynamic states of parcellations had better reproducibility than static parcellations using long acquisitions

We aimed to compare the performance of the Dypac algorithm to the performance of the k-Means algorithm. Dypac aggregated seed-based parcellations on short time windows (of about 3 minutes duration) while the k-Means used long time series (about 2.5 hours of resting state functional MRI signal). We compared the within-subject reproducibility of Dypac stability maps with the k-Means parcellations for three seed voxels associated with the PM-VIS, the dACC and the PCC subnetworks. Our results showed that most Dypac parcellations outperformed the k-Means parcellations (with long time series) in terms of reproducibility. Particularly, the reproducibility scores of the Dypac primary states outperformed the k-Means parcellations across seeds. For instance, the dACC seed and the Dypac primary states had a median Pearson correlation of 0.84 over a median correlation of 0.76 in the case of the k-Means parcellations. Similarly, the PCC seed and the Dypac primary states had a median correlation of 0.93 over a median correlation of 0.63 in the case of the k-Means parcellations (See Fig. 4). Also, Dypac secondary states had better reproducibility scores compared to the k-Means parcellations reproducibility in the cases of the dACC and the PCC seeds. For instance, the dACC seed and the Dypac secondary states had a median correlation of 0.79 over a median correlation of 0.76 in the case of the k-Means parcellations. Likewise, the PCC seed and the Dypac secondary states had a median correlation of 0.84 over a median correlation of 0.63 in the case of the k-Means parcellations (See Fig. 4).

**Figure 4.**
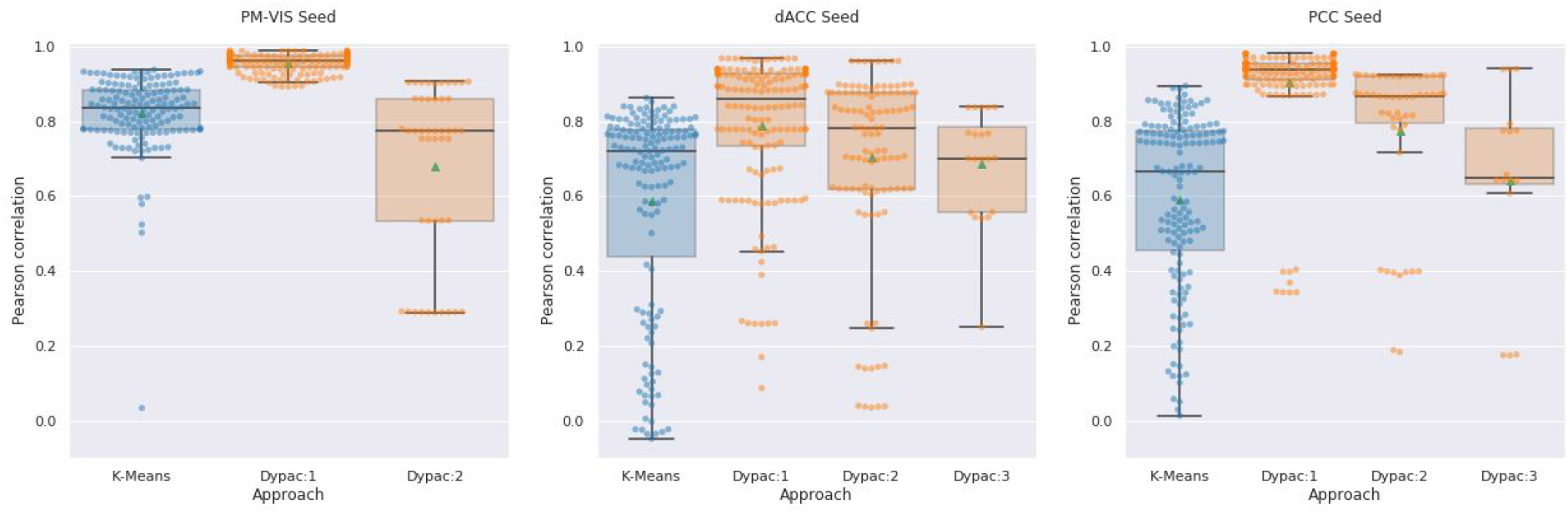
Dypac dynamic states of parcellations outperformed static parcellations based on long acquisitions in terms of within-subjects reproducibility. The within-subject reproducibility scores were computed between the two sets of five independent sessions. Both our Dypac parcellations and the k-Means parcellations used a total of ∼2.5 hours per set. Each algorithm was replicated 15 times per set with different seeds. The box plots represented the distribution of within-subject Pearson correlation scores. Dypac:1, Dypac:2 and Dypac:3 denoted the primary, secondary and third states of our Dypac algorithm. The green dots represented the mean Pearson correlation score for each distribution. We studied three seed voxels from the PM-VIS, the dACC and the PCC subnetworks. Ten subjects of the Midnight scan club dataset were investigated.

### 3.3 Visual evaluation of dynamic states of parcellations within- and between-subjects

To assess the reproducibility of parcellations, the Dypac dynamic parcellation method was applied on independent datasets for each subject, each dataset was composed of five sessions (∼for a total duration of 2.5 hours of data per subject) available in the MSC sample. We looked at the spatial reconfigurations of dynamic states of parcellations and tried to identify similarities and variations within-state (across replication sets, within-subject), across states (within-subject), and across subjects. We also added an extension for our method to consider spatially contiguous regions, which lead to similar conclusions as distributed parcellations (see Supplementary Material 2).

At the within-state level, we observed a high consistency between the dynamic states of the two replication sets, for all seeds and subjects. For instance, in subject MSC04 and the PM-VIS seed, the primary state maps showed high consistency in the left anterior insula (AI) region (Fig. 5, X=−38) and the supragenual anterior cingulate cortex (sACC) region (Fig. 5, X=−4). Similarly, the secondary state map of subject MSC02 and the dACC seed had consistent dACC region (Fig. 6, X=−4) and left AI region (Fig. 6, X=−38). Finally, the primary and the tertiary state maps of subject MSC02 and the PCC seed, had respectively consistent TPJ (Fig. 7, X=−38) and MPFC (Fig. 7, X=2) across the two replication sets.

**Figure 5.**
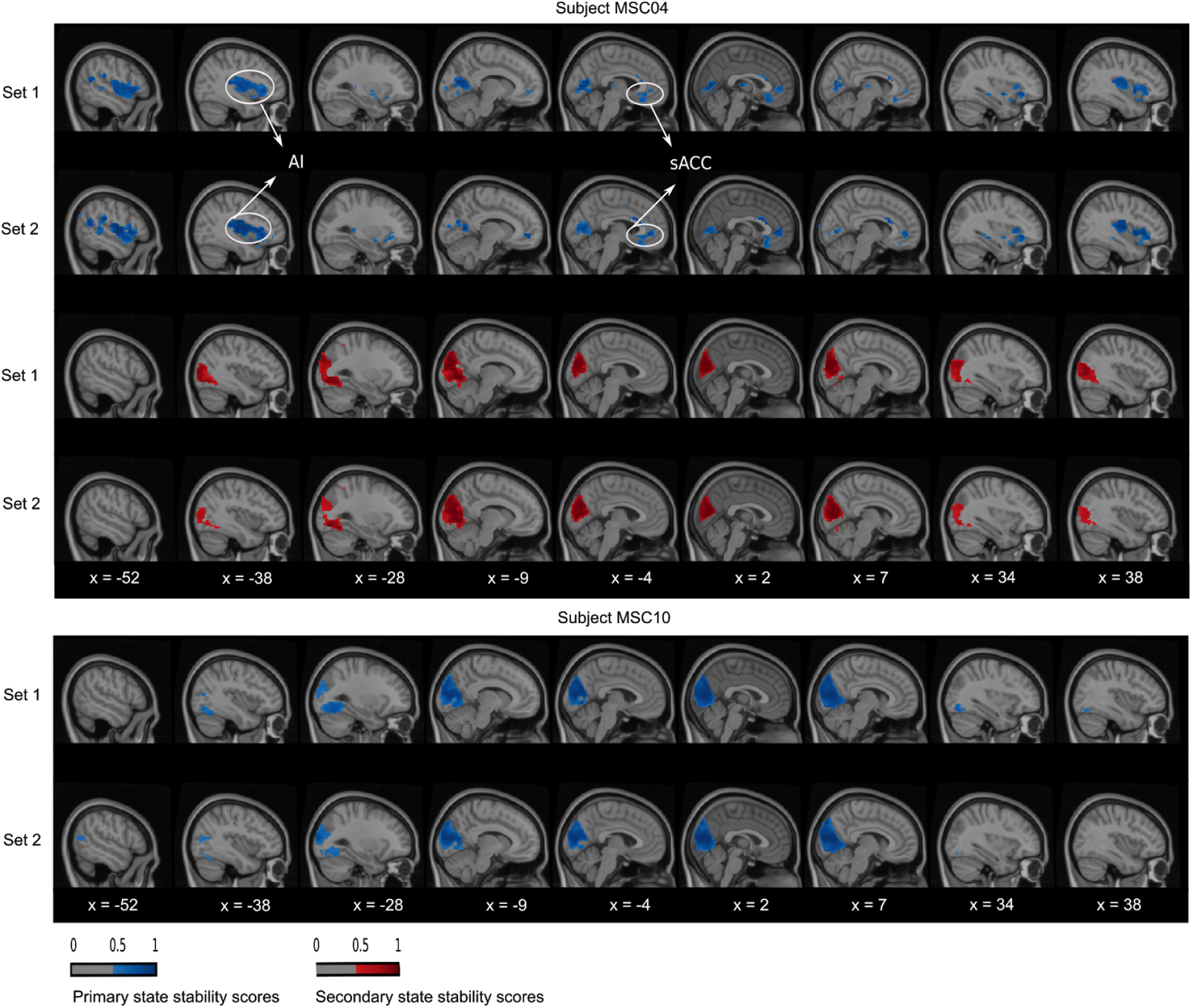
PM-VIS dynamic state stability maps were similar across sets at the within-subject level and variable across states and subjects. A complete matching between states was applied using the Hungarian method by maximizing the Pearson correlation between maps. The primary and secondary states were represented in, respectively, blue and red colors. A threshold was applied to keep only stability scores over 0.5.

**Figure 6.**
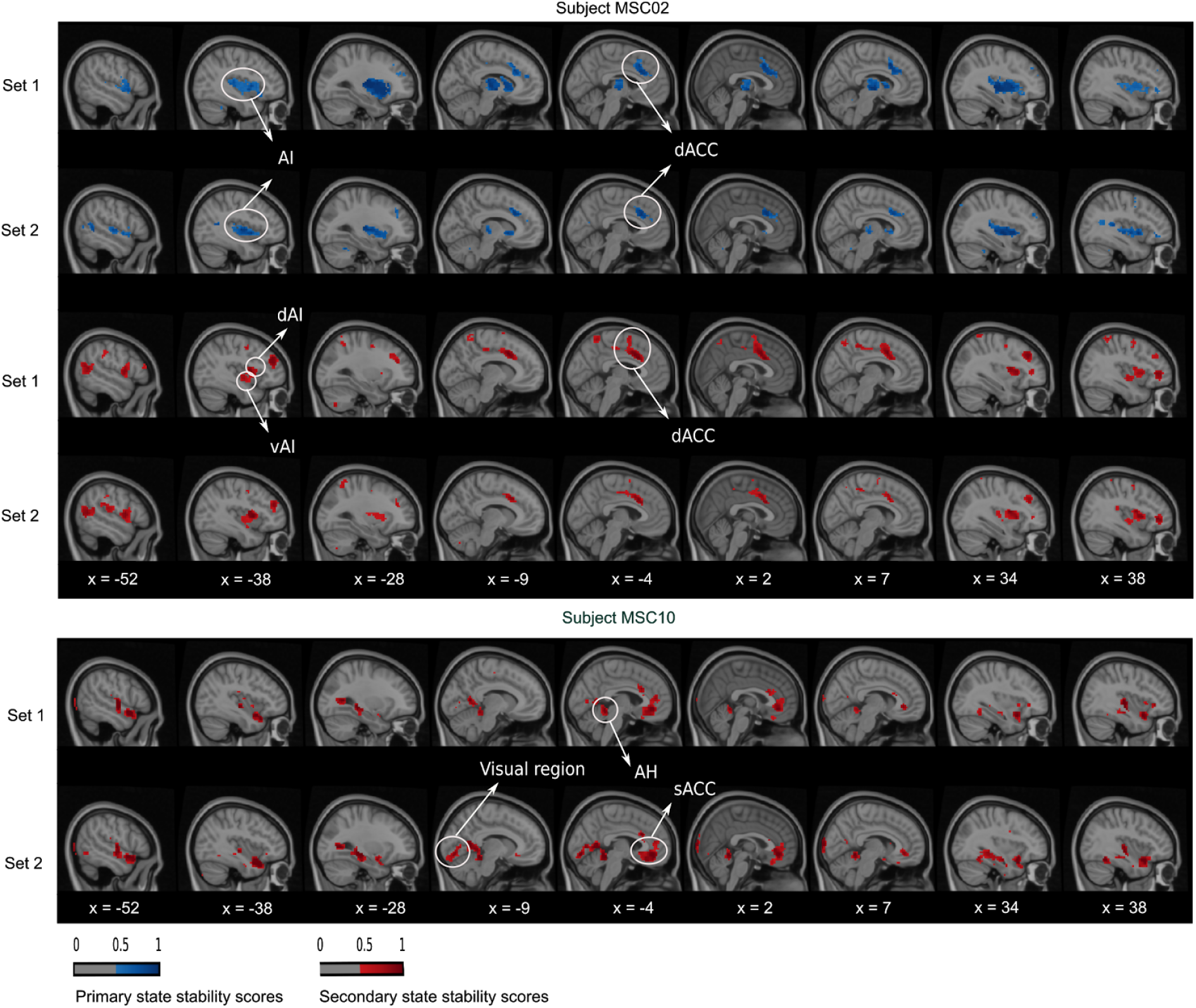
dACC dynamic state stability maps were similar across sets at the within-subject level and variable across states and subjects. A complete matching between states was applied using the Hungarian method by maximizing the Pearson correlation between pairwise maps. The primary and secondary states were represented in, respectively, blue and red colors. A threshold was applied to keep only stability scores over 0.5.

**Figure 7.**
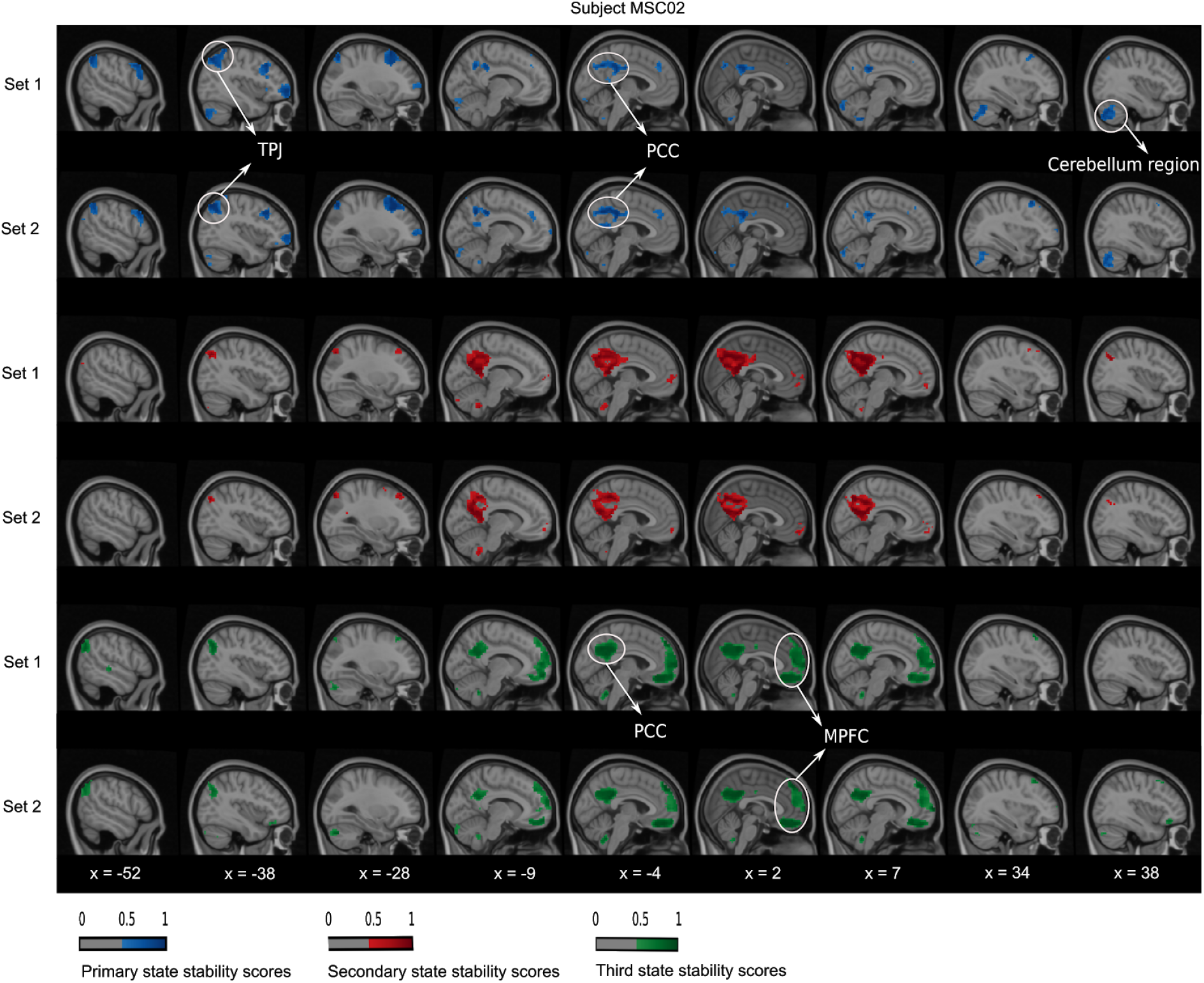
PCC dynamic state stability maps were similar across sets at the within-subject level. The primary secondary and third states were represented in, respectively blue, red and green colors. A threshold was applied to keep only stability scores over 0.5.

Across states, there existed different spatial reconfigurations within the same subject, especially in the cases of the dACC and PCC seeds. We observed some differences locally, at the level of a region surrounding the seed. For example, the region around the dACC seed was circumscribed and anterior in the primary state of subject MSC02, while the region shifted to the posterior direction in the secondary state (Fig. 6, X=−4). We also observed differences involving multiple regions distributed throughout the brain. Using again the example of subject MSC02 and the dACC seed, the entire AI was involved in the primary state, while the secondary state included only the dorsal anterior insula region (dAI) and the ventral anterior insula (vAI) regions (Fig. 6, X=−38). Similar local and distributed variations were observed with the PCC seed, which had up to three states in the case of subject MSC02 (Fig. 7). The primary state had a cortical region following the boundaries of PCC (Fig. 7, X=−4), along with distributed regions in the cerebellum (Fig. 7, X=38) and the left-temporoparietal junction (TPJ) (Fig. 7, X=−38). By contrast, the tertiary state included the PCC core (Fig. 7, X=−4) along the MPFC (X=2). Finally, the secondary state involved almost exclusively an extensive PCC region (Fig.7, X=−4).

At the inter-subject level, we found some overlapping in regions as well as completely inconsistent regions between subjects in their state spatial maps. As an example of overlapping regions, we found that the PM-VIS seed was characterized by highly similar visual cortex regions for all subjects (Fig. 5). Only subject MSC04 had an inconsistent primary state map that differed from other subjects maps. Here, we observed that the PM-VIS seed involved AI regions (Fig. 5, X=−38) and sACC regions (Fig. 5, X=−4). It was worth mentioning that no spatial matching was applied between state maps of two different subjects in figures 5, 6 and 7. Unlike the PM-VIS seed, the dACC and the PCC seeds had both overlapping and non-overlapping regions when we compared their state maps across subjects. As an example, we found some overlapping regions in the primary states of subjects MSC02 and MSC10 and the dACC seed (Fig. 6), e.g. overlapping AI region (Fig. 6, X=−38) and dACC region (Fig. 6, X=−4). For the same dACC seed, some subjects had non-overlapping regions such as the secondary states of subject MSC02 and subject MSC10 (Fig. 6). In the secondary state of subject MSC02, the dACC region (Fig. 6, X=−4) occurred as a dominant region along with AI regions (Fig. 6, X=−38). However, the secondary state of subject MSC10 was particularly characterized by the existence of the visual region (Fig. 6, X=−9), the anterior hippocampus (AH) and the sACC (Fig. 6, X=−4).

### 3.4 Dynamic states of parcellations are highly reproducible at the intra-subject level

We quantified the reproducibility of our dynamic states of parcellations at the within-subject level. First, we computed the within-subject consistency by means of a spatial similarity measure (i.e. Pearson correlation) between state stability maps associated with two sets of five sessions per subject. Each value in the stability map represented the probability of a voxel to belong to the cluster of a given seed. In these maps, we observed high within-subject reproducibility scores across states and seeds. For instance, most subjects had a reproducibility score that exceeded 0.8 in terms of the Pearson correlation across seeds in the cases of primary and secondary states. Although the third state rarely occurred, it had a high reproducibility score with more than 0.75 reproducibility score, e.g. for the dACC seed, both subjects MSC3 and MSC8 had, respectively, 0.78 and 0.75 Pearson correlation scores. Similarly, in the case of the PCC seed, subject MSC2 had a 0.9 correlation score. The PM-VIS seed was characterized by the highest reproducibility scores compared to the dACC and the PCC with more than 0.9. Except subject MSC04, all subjects had only one highly reproducible state (see Fig. 8). From one set, some states did not have a matched map from the second set and had, therefore a zero correlation score. For example, in the case of the PM-VIS seed and the primary state, subject MSC02 did not have a state in the replication sample that matched the primary state of the discovery sample (see Fig. 8).

**Figure 8.**
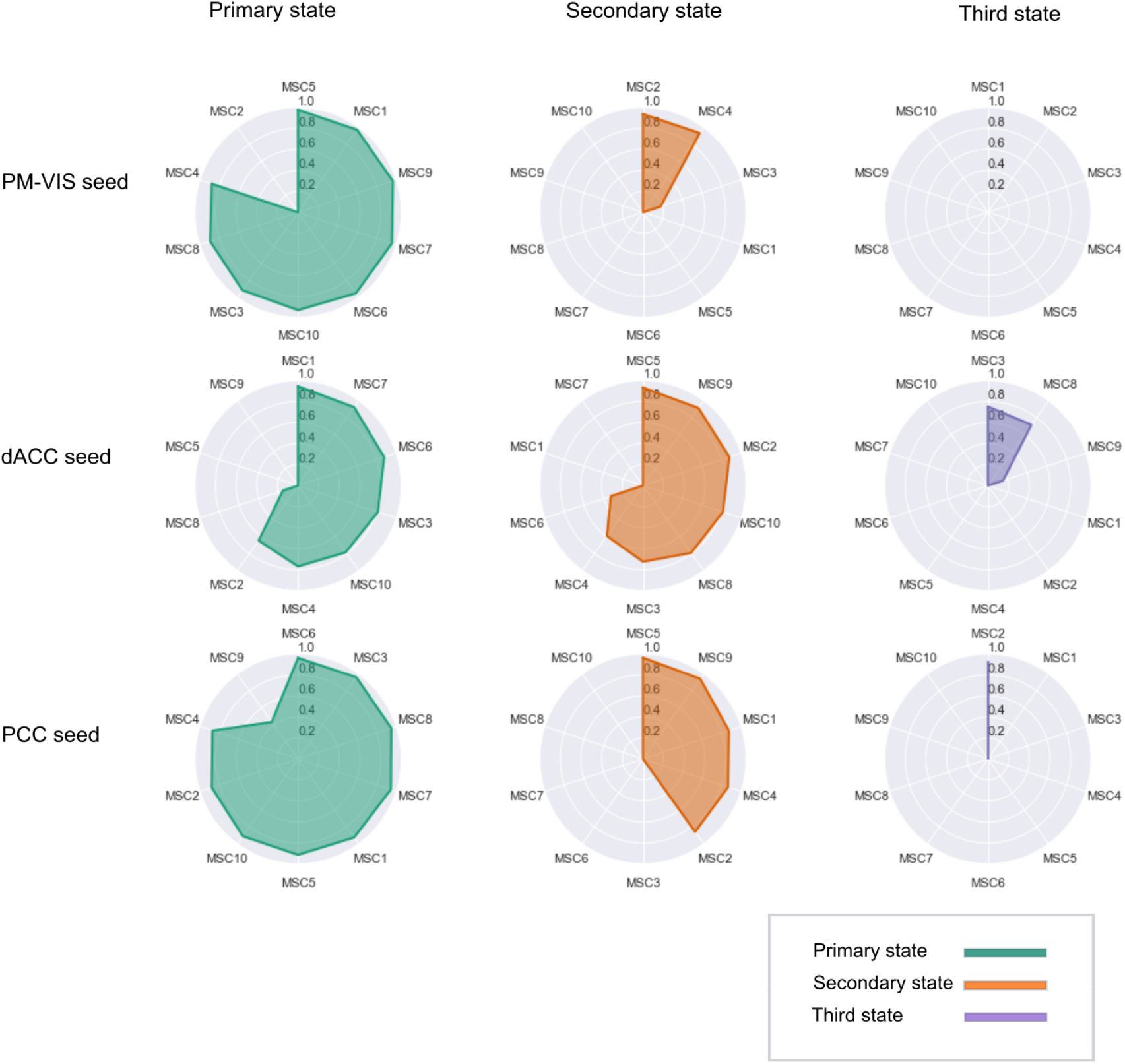
High spatial reproducibility of dynamic states of parcellations across subjects and seeds. Most subjects had two dynamic states of parcellations and the highest reproducibility score was found for the primary states of three subnetworks including the PM-VIS, the dACC and the PCC seeds. These reproducibility scores represented the similarity between state stability maps associated with two sets of data for each subject. Each set included five independent sessions. A complete matching between states was applied using the Hungarian method by maximizing the Pearson correlation between pairwise maps. States were sorted and labeled (i.e., primary state, secondary state, third state, etc.) based on their dwell time such that the primary state had the highest dwell time. Ten subjects of the MSC dataset were investigated, i.e. MSC1, MSC2, MSC3, etc.

### 3.5 Within-subject reproducibility of dynamic states of parcellations is substantially higher than between-subject reproducibility

In this section, our purpose was to contrast the dynamic states reproducibility within and between subjects. To this end, we cross-correlated their state stability maps as a measure of reproducibility and compared the results at the between- and within-subject levels, where the measures were derived from all the state maps simultaneously (i.e. pooling primary, secondary, etc). Our results showed that within-subject reproducibility scores outperformed the between-subject reproducibility scores with almost two disjoint distributions of correlation scores for all dynamic states and seeds, see Fig. 9. For example, the between-subject PCC-related scores did not exceed 0.78 while most within-subject reproducibility scores exceeded 0.8. Similar findings were observed in the case of the dACC. Only a few cases of the within-subject reproducibility scores fell within the distribution of between-subject reproducibility. We also investigated the impact of different parameters of our Dypac algorithm on the reproducibility of the dynamic states. We compared the results with different clusters; i.e. number of clusters in {12, 50} (see Supplementary Material 4.1), different window lengths in {30, 50, 100, 200} (see Supplementary Material 4.2), different cluster size thresholds in {5%, 10%, 20%} (see Supplementary Material 4.4), different smoothing kernels in {4mm, 6mm, 8mm} (see Supplementary Material 4.6), 15 different seed voxel coordinates from the visual network, the dACC and PCC subnetworks (see Supplementary Material 4.8) and, finally, different number of replications of seed-based parcellations with random seeds; i.e. number of replications in {1,5,30} (see Supplementary Material 4.9). Our conclusions on the reproducibility of the dynamic states of parcellations were valid for different parameters of the algorithm. That is, the within-subject reproducibility analysis robustly outperformed the inter-subject reproducibility across all ranges of parameters that were investigated. Moreover, differences in the distributions of within-subject reproducibility related to parameter changes were only subtle.

**Figure 9.**
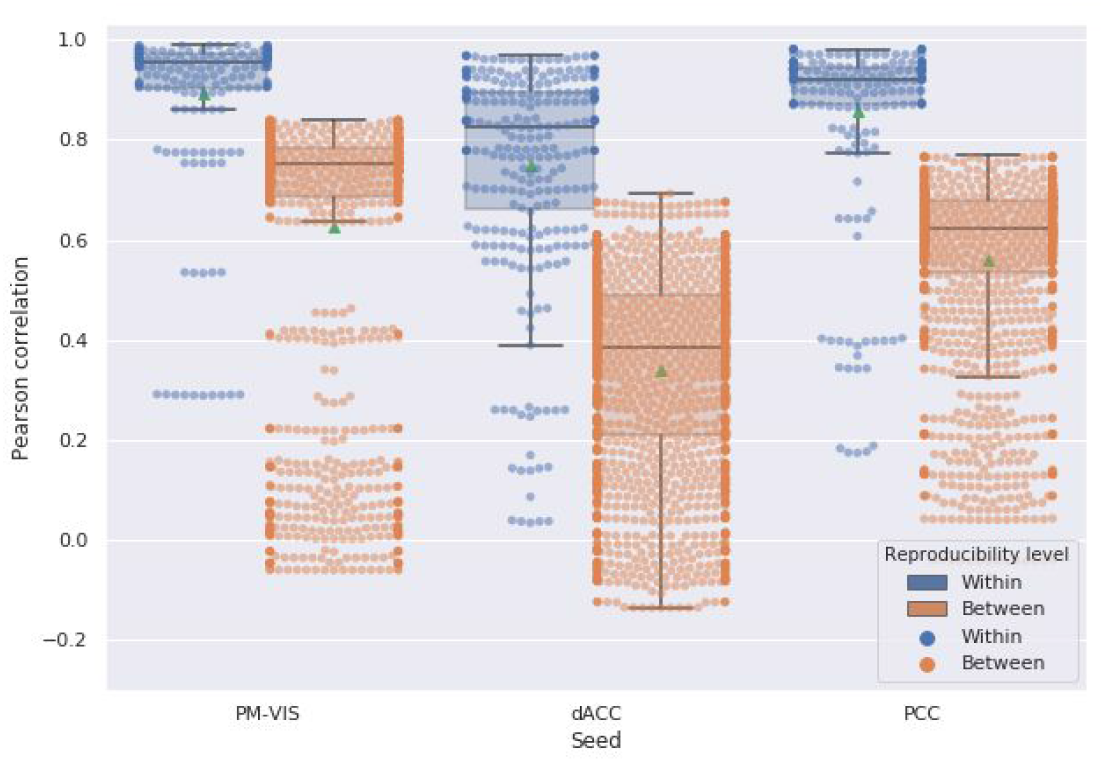
Within-subject reproducibility scores were higher than between-subject reproducibility scores for most dynamic states of parcellations. The Dypac algorithm was replicated 15 times with different seeds for each half of the data set. Number of replications of seed-based parcellations = 5. The green dots represented the mean of the Pearson correlation. We studied the PM-VIS, the dACC and the PCC seeds.

### 3.6 Dynamic state stability maps can reliably identify subjects

We evaluated the reliability of our dynamic state stability maps in identifying a particular subject among a pool of subjects using the fingerprinting experiment. Due to differences in the number of states per subject, we set up a fingerprinting by chance experiment as a baseline to verify the impact of these differences on the accuracy of the fingerprinting. We evaluated the accuracy of the fingerprinting by chance, and we compared it to the deterministic fingerprinting. Our results showed poor accuracy in the case of the fingerprinting by chance with an average accuracy score of 0.3 compared to the deterministic fingerprinting for which the average accuracy falled between 0.72 and 0.9 across seeds (See Fig. 10). These results showed a low impact of the differences in the number of states on the deterministic fingerprinting and, thus its accuracy scores were reliable.

**Figure 10.**
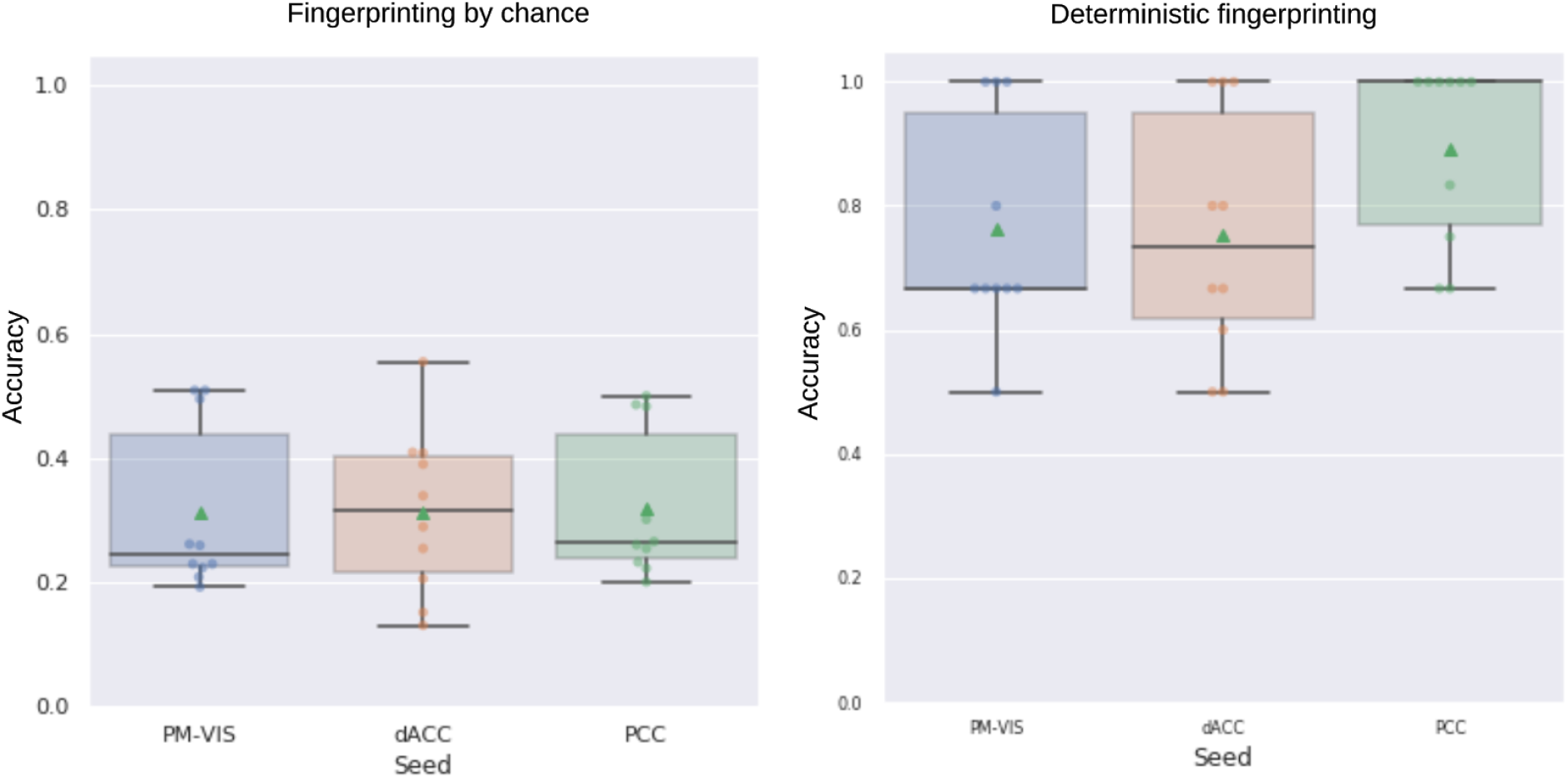
The deterministic fingerprinting had higher accuracy than the fingerprinting by chance. The accuracy of the fingerprinting represented the ratio of the successful fingerprinting over the total number of matchings both in the cases of the deterministic fingerprinting and the fingerprinting by chance. The state stability maps were generated from the split half sets of the Midnight scan club dataset and all these maps were pooled altogether in the fingerprinting. For a given map, we looked for the map that matched the closest map from the pool of all maps. In the case of a successful fingerprinting, the matched maps belonged to the same subject. The PM-VIS, dACC and PCC seeds were analyzed separately. Ten subjects of the Midnight scan club dataset were included.

The results of the deterministic fingerprinting experiment showed high accuracy results across seeds, with more than 0.72 average accuracy scores. This confirmed that many subjects were successfully fingerprinted based on one of their state maps. The highest accuracy results were associated with the PCC seed with an average accuracy of 0.9 across states. Also, the dACC state maps had a high average accuracy score of 0.72. Similarly, the PM-VIS had an average accuracy of 0.78. In the case of a failure, two state stability maps were highly correlated but their maps were not associated with the same subject. Most failures were associated with the PM-VIS and the dACC. Overall, these findings confirmed that our dynamic state stability maps were reliable in the delineation of subjects (See Fig. 10).

We further reported the Pearson correlation scores associated with the deterministic fingerprinting experiment. The distribution of correlation scores across subjects allowed us to quantify the spatial similarity across subjects. Most importantly, failed fingerprinting allowed us to have a better understanding of the degree to which state maps were similar across subjects. Our results showed that the successfully matched maps had high pearson correlation scores. For instance, the dACC and the PCC seeds had, respectively 0.8 and 0.9 Pearson correlation scores. In the case of failures, the lowest scores were associated with the PM-VIS seed with a 0.5 median correlation, while the highest scores were associated with the PCC seed with a 0.7 median correlation (See Fig. 11). The high Pearson correlation scores in the case of failures; correlation > 0.6, may be associated with spatially similar maps across subjects. Here, the PCC had the highest spatially similar state maps between subjects. Overall, the high accuracy and the high correlation measures confirmed the reliability of the fingerprinting in identifying a given subject based on his dynamic state map.

**Figure 11.**
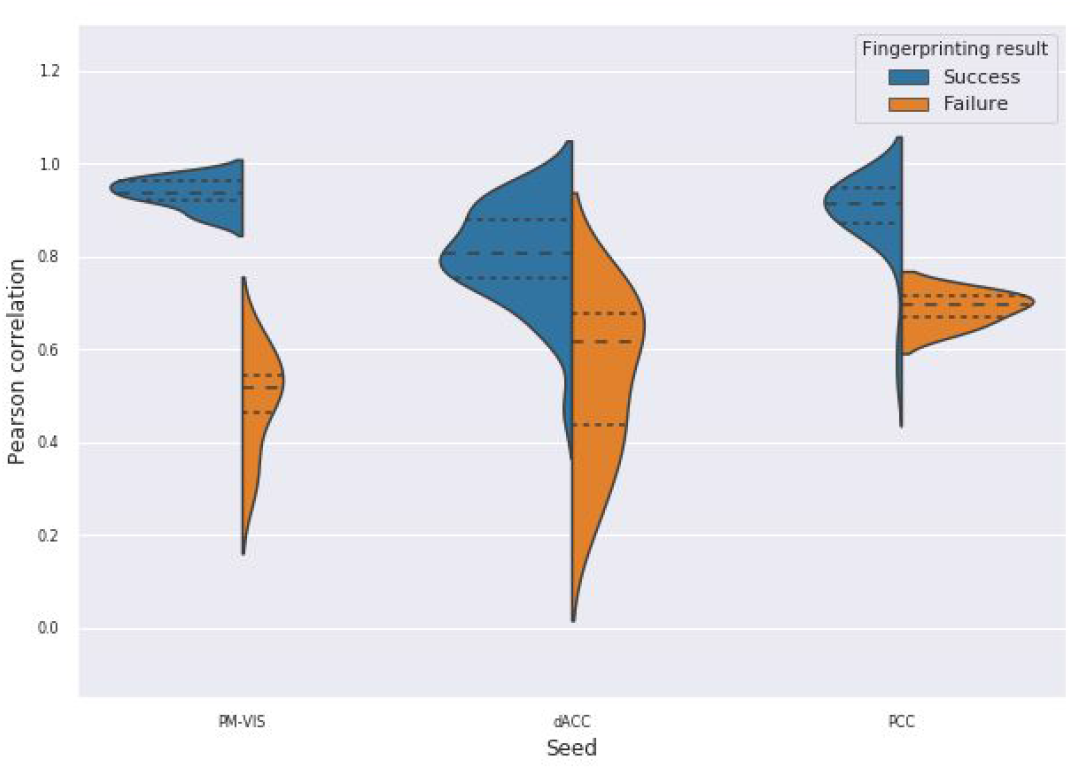
The fingerprinting experiment showed high reliability of state stability maps in delineating subjects across seeds. We showed the correlation scores for the deterministic fingerprinting experiment results. The spatial similarity was computed between pairs of state stability maps in terms of Pearson correlation. If the correlated maps were associated with the same subject, it was considered a success fingerprinting (blue color). Otherwise, the correlated maps were associated with different subjects. This was considered a failure (orange color). We studied three seeds from three subnetworks including the PM-VIS, the dACC and the PCC. Ten subjects of the Midnight scan club dataset were included.

**Figure 12.**
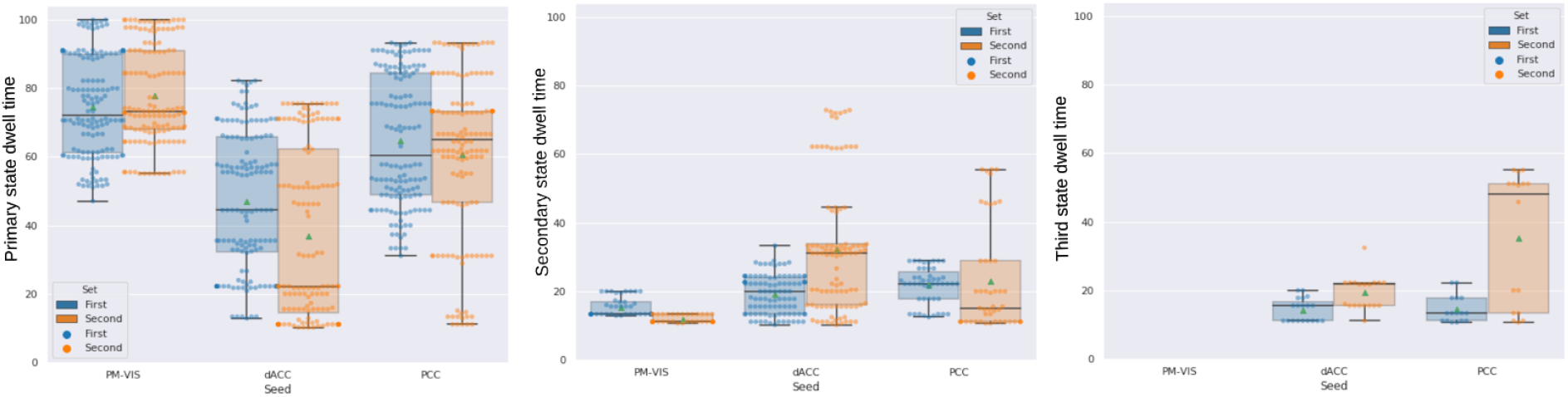
Dwell time of dynamic states of parcellations was inconsistent across the two sets of independent data in the cases of the dACC and PCC seeds. The dwell time across states was computed by summing durations of time-windows per state from two sets of five independent sessions for each subject. Three seeds were investigated including the PM-VIS, the dACC and the PCC. The seed-based parcellations number = 5. The number of replications of the Dypac algorithm = 30. The seed-based parcellations were clustered into 12 clusters. The smoothing kernel = 6 mm, the cluster size threshold = 10%, the Dice threshold = 0.3. Number of timepoints in the window length = 100. All the ten subjects of the Midnight scan club dataset were included. Green dots represented the mean of the dwell time value.

### 3.7 Dynamic state dwell times were not reproducible across replications for the dACC and the PCC seeds

We aimed to get a better understanding of the dwell time reproducibility over time. To this end, we performed a spatial matching of states between two sets of independent sessions in terms of the Pearson correlation. This matching was based on the Hungarian method. Therefore, only the dwell times associated with spatially reproducible states were included. Our results showed an inconsistency between states dwell time for most states between the two sets in the cases of the dACC and the PCC seeds. For instance, the primary state and the dACC seed had 43% median dwell times for the first set over 22% median dwell times for the second set. Similarly, the PCC and the secondary states had 23% median dwell times for the first set over 17% median dwell times for the second set. Unlike the dACC and the PCC seeds, the dwell times of the PM-VIS had higher reproducibility across the two sets. For instance, the primary state and the PM-VIS seed had 62% dwell times for the first set over 64% dwell times for the second set (See Fig. 12). Overall, the dynamic states dwell times were not reproducible across the two sets of independent data for the dACC and the PCC subnetworks. The PM-VIS showed higher levels of consistency between dwell times of the two sets.

We also observed that some states might have a high dwell time but no matching in the second set. For instance, in the case of subject MSC09 and the dACC seed, there existed five states with the following dwell times for the first set: 31.11%, 12%, 11.11%, 10.66%, 10.22%. However, this subject had only two states in the second set with the following dwell times 22.22% and 13.33%. After the spatial matching, the matched primary states had 11.11% and 22.22% dwell times for the first and second set, while the secondary state had 10.22% and 13.33% for the first and second sets, respectively.

## 4. Discussion

In this paper, our overall objective was the identification of dynamic states of brain parcellations in individual resting state fMRI data. Our first main finding was the existence of highly similar spatial parcellations extracted from short time windows, sometimes separated by several days. This led us to propose a dynamic cluster analysis to extract dynamic states of parcellations. These dynamic states were markedly different in terms of the brain regions involved, despite being derived from the same seed region and the same subject. We also found that dynamic states of parcellations were subject-specific, highly reproducible, and are reliable enough to successfully differentiate subjects in a fingerprinting experiment with high accuracy.

In the literature of brain parcellation, the only approach that had consistent findings with our approach was published by Salehi and colleagues (Salehi et al., 2020; Salehi, Greene, et al., 2018; Salehi, Karbasi, et al., 2018). We both suggested an approach that contradicts the notion of a fixed functional parcellation of the brain. The main difference with our work is that Salehi and colleagues generated a different parcellation for different cognitive states in a series of task datasets (i.e. motor task, working memory, rest, etc.), while in our approach we identified different dynamic states of parcellations in short windows of a single cognitive state (resting state). Our results on dynamic brain parcellation is in line with several studies which showed that brain connectivity is highly dynamic, with recurring spatiotemporal patterns of brain subnetworks (Abrol et al., 2017; Allen et al., 2014; Vince D. Calhoun et al., 2014; Donnelly-Kehoe et al., 2019; Hutchison et al., 2013; Korhonen et al., 2017) both in the case of spatially distributed regions or even in the case of spatially contiguous regions (see Supplementary Material 2). For instance, Iraji and colleagues demonstrated the existence of spatial fluid interactions between intra- and inter-networks relationships, emphasizing the dynamic interplay between segregation and integration (Iraji et al. 2018). Researchers raised the need for new computational methods to reveal robust, interpretable reconfigurations in the complex and high-dimensional feature space of dynamic fMRI data (Chen et al., 2017). Such methods would allow a better understanding of the individual differences in internal state changes over short time scales (Chen et al., 2017). In the brain parcellation literature, our approach is, to our knowledge, the first attempt to shed light on this dynamic brain organization at fine temporal scale (in the order of few minutes) and at a voxel level (i.e. without reducing the dimensionality in a fine-grain parcellation).

To evaluate the quality of our dynamic states of parcellations, we relied mainly on a reproducibility analysis. We found that our reproducibility scores markedly outperformed static brain parcellation scores (Kong et al., n.d.). For instance, the reproducibility scores observed in the case of our visual dynamic states of parcellations were near perfect (with an average correlation scores of 0.95), while static visual parcels using about 2.5 hours of data had 0.85 correlation score in average (See Fig. 4). Moreover, visual static brain parcels reported previously in the literature did not reach near perfect similarity scores (with an average Dice score of 0.85) (Kong et al., n.d.). Such high reproducibility scores were also observed for all three seeds such that dynamic states scores outperformed static parcels scores (See Fig. 4). Another important consideration was that parcels reproducibility depended on the spatial location in the brain in the case of static parcellations, while we observed the same ranges of values for the three seeds. For instance, static parcellations reproducibility ranged between 0.7 and 0.85 average Dice scores in the cases of temporal and visual cortices regions, respectively (Kong et al., n.d.). Interestingly, our spatial correlation similarity measure exceeded 0.85 for the three seeds for the primary states, and exceeded 0.7 correlation score for secondary and third states for almost all subjects and seeds (See Fig. 4). Recent attempts towards better reproducibility scores of static connectivity measures relied on long fMRI acquisitions (Gordon, Laumann, Gilmore, et al., 2017). Even though similarity measures were improved (with 10 minutes and 50 minutes of data, authors got, respectively 0.6 and 0.7 average Dice scores), these measures plateaued after ∼40 minutes of acquired signal with a maximum average of 0.7 Dice score across the ten subjects of MSC dataset. Using long acquisitions of functional MRI data, the comparison of the k-Means parcellations with our dynamic states confirmed that a dynamic approach of the parcellation problem resulted in improved reproducibility (See Fig. 4). Moreover, higher Dice scores were observed for pairs of seed-based parcellations as extracted from a few minutes (∼3 minutes) of fMRI data within a given dynamic state (see Supplementary Material 3). Overall, many studies have aimed to derive static brain parcellation approaches either at the group or the individual level (See (Arslan et al. 2017) and (Eickhoff et al. 2018) for a review) and yet, no technique had highly reproducible individual parcels. This leads to the conclusion that human brain parcellation is a hard ill-posed problem (Arslan et al., 2017). Our results demonstrated that the main limitation associated with static brain parcellation approaches is neither the quality of employed clustering algorithms, the quality of fMRI data in specific brain regions nor the duration of fMRI acquisitions. The main issue was the incorrect formalization of functional brain parcellation as a static problem, which did not take into consideration the dynamic organization of the brain. Specifically, we showed here that a very basic clustering algorithm, k-Means, leads to highly reproducible parcellation maps when applied on short fMRI time series (a few minutes) with a dynamic approach.

Another important consideration was the comparison of the reproducibility scores within and between subjects. Previous studies reported that within-subject similarity of dynamic states of parcellations was substantially higher than inter-subject similarity (Kong et al.). Our quantitative and qualitative evaluation were consistent with these findings (See Fig. 4, 5, 6, 7 and Supplementary Material 4) and we suggest that dynamic states of parcellations captured some of the variability between subjects. To further evaluate the reliability of these dynamic state stability maps in identifying a particular subject from a pool of ten subjects from the Midnight scan club dataset, we implemented a fingerprinting experiment. Even though the fingerprinting failed in a few samples, results on the ten subjects of the MSC dataset showed high scores with 0.6 and 0.7 average accuracy scores for the PCC and the dACC seeds, respectively (See Fig. 11). Due to the small sample size, in the MSC dataset, investigating and validating our results on larger samples in the future, needs to be considered. Researchers already demonstrated the variability in functional connectivity profiles as a reliable fingerprinting to identify subjects from a large group (Finn et al., 2015). Our state stability maps were derived from binary cluster maps that eliminated a huge amount of the fine details present in a connectivity map. Despite such dramatic dimensionality reduction in dynamic state maps, it preserved enough relevant information to reliably delineate subjects, especially in the case of highly cognitive networks (i.e., the PCC subnetwork). Our PCC and PM-VIS accuracy scores actually outperformed the accuracy scores of connectivity maps-based fingerprinting in some networks (DMN and the salience), as reported in (Badhwar et al., n.d.), but this observation may also reflect the fact that we used much longer individual fMRI time series and less subjects.

In the context of our Dypac algorithm, we showed that the reproducibility of the dynamic states of parcellations were robust to the choice of different parameters (See Supplementary Materials 4). Still, it is important to mention that the number of clusters k is a critical parameter, and can be used to uncover the pseudo-hierarchy of brain subnetworks. Here, we simply checked that “states” could be identified at two different resolutions (i.e., 12 and 50), but it remains to be tested how the number of states vary with resolution, and whether dynamic parcellation follows a pseudo-hierarchical organization as was previously described by static parcellations (Urchs et al., 2017).

Added to the spatial reproducibility analysis, our temporal analysis showed that the dwell times of dynamic parcellations were inconsistent across the two sets of independent data. This may either indicate algorithmic variability in the estimation of dwell times, maybe linked to our choice of threshold on inter-parcel similarity to define states, or physiological variability where a given subject expresses markedly different states over long time scales. We believe the latter to be more plausible, but further validation of this hypothesis would require data which directly manipulate cognitive states across replication states, e.g. using tasks, and is outside of the scope of the present paper. We note however, that the high variability of dwell times across replication sets is probably the factor that drives the “glass ceiling” in reproducibility of static methods: even the definition of what is the primary state can change over long time scales, so averaging across states is not sufficient to stabilize parcel estimates.

Another important aspect of the evaluation of the dynamic states of parcellations was their neurobiological validity. In the absence of brain organization ground truth, current parcellation work capitalized on replication, robustness and convergence as criterias for biological validity. However, the observed variations in the shape and position in the spatial patterns across individuals suggest that these patterns are likely to be associated with physiological or cognitive processes (Eickhoff et al. 2018). As pointed out in the previous section, our quantitative results support the neurobiological validity of our dynamic states of parcellations. Qualitatively, we observed that many regions from the dynamic states of parcellations relate to previous literature when studying subnetwork dynamics. For instance, the regions of the dACC state stability maps overlapped with the salience network regions as reported by (V. Menon, 2015) including the insula and the anterior cingulate cortex (i.e., sACC). Consistently with previous research, we frequently observed the AI and the dACC either in the primary or the secondary states of parcellations. These regions were among the most frequently activated regions in functional neuroimaging research (Buchsbaum et al. 2005; Yarkoni et al. 2011). In some dynamic states of parcellations, we observed a high stability around the motor and premotor regions of the dACC maps (See Fig. 4). This may be explained by the existence of a functional coupling between the AI and the dACC that facilitates a rapid access to the motor system (V. Menon, 2015). Similarly to the dACC seed, we also observed that the PCC is multistate. In the literature, researchers observed a high spatial heterogeneity in the PCC (Leech & Sharp, 2014; Margulies et al., 2016), but in our case dynamic states were observed from a single seed and subject. Dynamic states of parcellation of the PCC seed identified, mainly ventromedial prefrontal cortex regions (i.e., including the MPFC), the superior parietal cortex regions and the precuneus. Leech and Sharp surveyed the different studies that investigated the variations in PCC activity with arousal state and its interactions with other brain networks. Authors suggested that the high heterogeneity of the PCC activity was attributed to its important role in regulating the balance between internal and external attentional focus (Leech & Sharp, 2014). While higher order seeds revealed multistate maps including the dACC and the PCC seeds, our results showed the PM-VIS was also multistate for some subjects (e.g. subject MSC04) even though most subjects had a monostate PM-VIS subnetwork. Consistently to resting state functional connectivity studies, the visual cortex was considered as a unimodal system since it had a maximal distance along the principal gradient between the visual and the DMN which was considered as a highly heteromodal network (Margulies et al., 2016). Overall, these qualitative observations support our hypothesis that dynamic states are driven by biological validity rather than methodological effects (Eickhoff et al., 2018).

In addition to biological meaningful brain parcels, researchers hypothesized the existence of some non-meaningful parcels that may occur due to physiological sources or other non-neural effects such as head motion (Chen et al., 2017; Leech & Sharp, 2014). For the scope of this paper, we did not characterize these sources and we consider this an important follow up question to be studied for more seed regions. However, with the proposed method, a large number of time windows were not associated with a state, if no robust parcel configuration was identified. This feature of the method may help to mitigate the influence of confounding effects on dynamic brain parcellations. However, sources of physiological noise with highly consistent spatial distribution, such as cardiac noise and motion artifacts, may still lead to robust spatial parcellation states. We also showed there was no sessions effect on the identified dynamic states (See Supplementary Material 5).

The main conclusion of this work is that stable brain parcellations emerge from a dynamic analysis considering short time windows, which challenges the notion of a fixed, static brain parcellation estimated from very long time series (Gordon, Laumann, Gilmore, et al., 2017). But this observation was restricted to a few seed regions in the brain, and an important point of discussion is whether the Dypac algorithm could be generalized to the full brain ? The core generation of brain parcellations was a simple k-Means algorithm applied on full brain data, and we trust that our conclusions extend beyond the handful of seeds which we considered. We notably confirmed that our conclusions generalize to many neighboring voxels around selected seed regions, see Supplementary Material 4.8. One way of conceptualizing k-Means as a sparse spatial decomposition: each brain voxel is associated with only one brain parcel, which naturally leads to parcels (or seed-based parcellation) that include only a small portion of the brain (for a large number of clusters k). Similarly, our dynamic analysis can be conceptualized as a sparse temporal decomposition: for each brain voxel, only a subset of time points are associated with a single brain parcellation (or seed-based parcellation), and many time points are associated with no brain parcellations at all. The Dypac algorithm is, thus a double sparse space-time decomposition technique, which focuses only on temporally recurring and highly spatially similar brain parcels. Dypac could, in theory, be applied on the brain parcels associated with all brain voxels simultaneously, represented as a basis of one-hot spatial encoding vectors, although the memory cost of the hierarchical clustering step would become prohibitive. We are working on a modified Dypac algorithm using k-Means both for parcellations generation and aggregation, which scales to the full brain^8^ even at high spatial and temporal resolutions. We would like to emphasize at this stage that the idea of applying a space-time decomposition to fMRI data is old, at least as old as spatial independent component analysis, ICA (McKeown & Sejnowski, 1998) which identified brain networks as a temporal mixture of spatially independent components, including noise. This approach was extensively applied for more than two decades in fMRI research (see survey of (Beckmann, 2012)). Some more recent space-time decomposition of fMRI data explicitly included a spatial sparsity constraint (Dadi et al., 2020). Based on a visual comparison of our dynamic states of parcellations and ICA spatial maps, we identified overlapping patterns especially in the case of the visual and the default mode network including the PCC/precuneus, the MPFC regions and regions of the dorsal attentional subnetwork (Beckmann et al., 2005; Damoiseaux et al., 2006; Zuo et al., 2010). A major limitation of the ICA technique was the variability of functional resting state signal at the individual level added to its random initialization ((Beckmann et al., 2005; Damoiseaux et al., 2006; Zuo et al., 2010)). An intriguing possibility is that dynamic states of parcellations would converge towards some similar spatial patterns as those identified by ICA, although with superior spatial stability. This possibility will need to be further investigated. Such observation would create a bridge between traditional cluster analysis and space-time decomposition techniques such as ICA, even though the underlying formalism is quite different.

The existence of dynamic states of parcellation could have important implications for graph-based analysis of brain networks. In such circumstances, building brain graphs using these parcels remains a challenging question. As discussed in the preceding paragraph, a full-brain extension of Dypac would, in practice, be a new flavor of space-time decomposition of fMRI data which may result in improved characterization of brain graphs compared to either static clustering-based parcels or ICA techniques. Unlike traditional static brain parcellation, and like an ICA, our dynamic states of parcellations have spatial overlap (if they associate with different states of the same voxel). Such types of decompositions are straightforward to apply in a graph-based analysis, as indeed many graph analysis of fMRI data have relied on ICA decomposition to define nodes (Yu et al., 2017). The key difference between static parcellations and space-time decomposition is how fMRI data are embedded in the parcels. For static parcellation, the embedding is univariate in nature, as each parcel is treated separately: an average time series is generated for each parcel, or a principal component analysis is applied to the time series inside the parcel (Eickhoff et al., 2018). For space-time decomposition, the embedding is multivariate in nature, as the parcels (or spatial modes of decomposition) are treated jointly using a multivariate regression analysis to fit a full brain activity volume at each time point (V. D. Calhoun et al., 2001). When applied to traditional static parcellations generated by a cluster analysis, this regression step is equivalent to extracting the average time series per parcel. However, in the presence of overlap between brain parcels the regression solution is different. In this paper, the implementation of Dypac is a proof of concept and it is restricted to a seed brain region, thus, it is not directly applicable to generate full brain graphs. As mentioned earlier, we are working on a full brain extension of Dypac. In this case, we intend in the future to assess the practical advantages of Dypac to study fMRI graph analysis.

Before Dypac is applied to graph analysis, it will notably be important to evaluate whether dynamic parcels capture fMRI dynamics with greater fidelity than static parcels ? In the case of static brain parcels, researchers often investigated two types of quality metrics (Thirion et al., 2014): first, are the parcels spatially reproducible (the focus of this paper) ? and, second, are the parcels homogeneous? Parcel homogeneity means that the time series of voxels inside a parcel are similar (i.e., highly correlated to each other). Generalizing the concept of homogeneity to space-time decomposition implies to embed fMRI activity in a set of spatial modes, as explained in the section above. It is then possible to compare raw voxel-based representation of brain activity, or brain maps, against a compressed version using parcel-based information only (Craddock et al., 2012; Yu et al., 2017). The quality of this compression is the space-time decomposition analogue of parcel homogeneity. Because we did not apply Dypac to the full brain in this work, we were not able to directly generate embeddings and test whether the increase in spatial reproducibility of brain parcels also came with an increase in compression quality. Up to date, there are only few guidelines towards the application of ICA maps or brain atlases as a better choice to reduce the dimensionality of brain networks in neuroimaging studies (Dadi et al., 2019), although one recent work already shows a promising result in that regard: decomposition into overlapping, smooth spatial modes lead to much improved compression of statistical brain maps (Craddock et al., 2012; Yu et al., 2017). An important area of future work is to extend dynamic brain parcellation to the full brain, rather than seed voxels, and test whether the compression of raw brain activity is improved over static brain parcels.

## Conclusion

To summarize, we show that a simple clustering technique such as k-Means can lead to highly reproducible functional parcellation associated with a particular brain seed, even when applied on short fMRI time series (a few minutes). We proposed a method to identify the main states of these dynamic parcellations, and showed that their spatial distributions were subject specific. The main limitation of previous work on functional brain parcellation may be due to the over simplified static parcellation approaches. This limitation may be biasing all neuroimaging analyses that rely on static parcellation as a dimensionality reduction step, including many graph-based neuroimaging analyses. Therefore, we urge the neuroimaging scientific community to replace static brain parcellations by dynamic parcellation approaches, in order to properly capture the rich interactions between brain subnetworks. Dynamic parcellations may thus impact widely applications of brain connectivity in health and disease.

## Acknowledgements

The authors are grateful to Dr Catie Chang for providing precious feedback during early iterations of this project and to the Midnight scan club (Gordon, Laumann, Gilmore, et al., 2017) for making MSC dataset available for our analysis. This work was funded by “The Courtois project on neuronal modelling”, based on a donation of the Courtois foundation, and funds of the Natural Sciences and Engineering Research Council (NSERC) to PB. YZ is a research fellow of the “Institut pour la valorisation des données” (IVADO). PB is a research fellow (Chercheur Boursier Junior 2) of the “Fonds de Recherche du Québec - Santé” (FRQ-S).

## Footnotes

1 https://hub.docker.com/r/simexp/niak-cog/

2 http://niak.simexp-lab.org/pipe_preprocessing.html

3 https://docs.python.org/3.4/library/multiprocessing.html?highlight=process

4 https://scikit-learn.org/stable/modules/generated/sklearn.cluster.KMeans.html

5 https://nilearn.github.io/modules/generated/nilearn.regions.connected_label_regions.html

6 https://github.com/SIMEXP/dynamic-states-parcellations

7 https://identifiers.org/neurovault.collection:6642

8 https://github.com/courtois-neuromod/dypac

