## Supplementary Materials for "Unraveling reproducible dynamic states of individual brain functional parcellation"

### Supplement 1

#### Whole brain parcellations had no clear structured states

We aimed to test the existence of structured states within the whole brain reconfiguration. To this end, we proposed a whole brain parcellation approach across sliding windows. First, we generated several whole brain k-Means parcellations across sliding windows. Then, we computed the adjusted rand index[1] similarity matrix between all pairs of sliding window parcellations. Our results showed very low similarity scores between pairs of k-Means parcellations and no clear structure was detected from the whole brain parcellation. For instance, the adjusted rand index similarity matrix for subject MSC01 did not show any groups of homogeneous sliding window parcellations (See Fig. 1-A). Also, the distribution of the similarity scores between sliding window parcellations were very low; i.e. adjusted rand index scores < 0.01 (See Fig. 1-B). This clearly showed that, although individual brain subnetworks follow reproducible “states”, there was no strong coupling across different brain subnetworks which would lead to full-brain parcellation states.

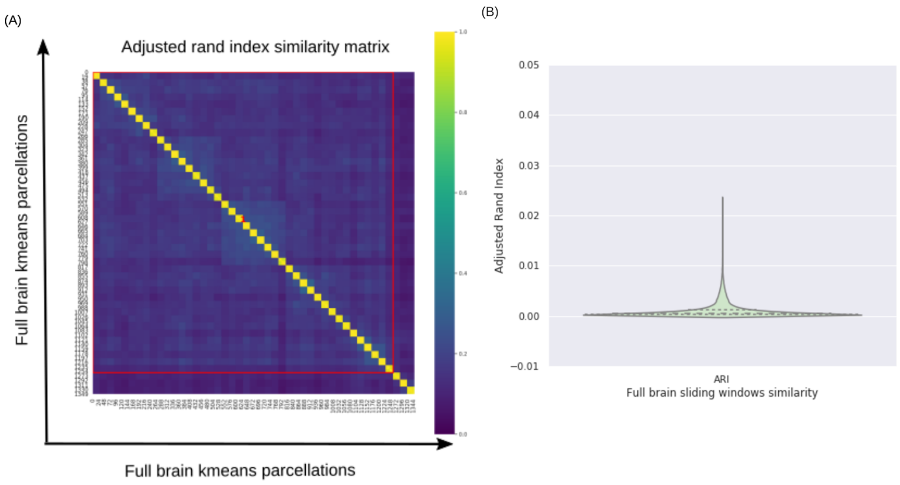


**Figure 1.** (A) The similarity matrix of the full brain parcellations were dissimilar across sliding windows in the case of subject MSC01. (B) Distribution of the adjusted rand index scores across subjects. The number of timepoints in the window length = 100. The number of sliding window replications = 30. A total of 1349 brain parcellations were included across five sessions. Three subjects of the Midnight scan club dataset were included (MSC01, MSC02, MSC03).

### Supplement 2

#### Dynamic states can be identified for spatially contiguous regions

Our dynamic states of parcellations were generated by aggregating several seed-based parcellations distributed over the brain. Many parcellation algorithms proposed in the past have enforced the parcels to be spatially contiguous. In order to make our method more

[1] https://scikit-learn.org/stable/modules/generated/sklearn.metrics.adjusted_rand_score.html

directly comparable to these prior works, we also implemented an extension in which each distributed parcel was subdivided into spatially contiguous regions. To this end, we extracted the regions having a minimum of 50 voxels. For the dACC and PCC seed voxels, we reported, side-by-side, both the state stability maps associated with the spatially contiguous regions and the distributed regions throughout the brain (see Fig. 2 and Fig. 3). Our results showed a good consistency between the distributed spatial parcels (or clusters) surrounding the seed and the spatially contiguous regions. For instance, the dACC primary state and subject MSC01 had a high overlap between the dACC distributed parcel and the dACC contiguous regions (See Fig. 2, X=4). Also, the distributed dACC parcel of the secondary state and subject MSC10 had a high overlap with the spatially contiguous dACC region of the primary state (See Fig. 2, X=4). In the case of the PCC, the primary states of subject MSC01 were highly overlapping for both the spatially distributed parcel surrounding the seed and the contiguous region (See Fig. 3, X=4).

The aggregation of seed-based contiguous regions suppressed the distributed regions, by construction. For instance, in the primary state and subject MSC10, the sACC region only occured in the state map of the spatially distributed regions (See Fig. 2, X=4). Also, in the case of subject MSC10 and the third state of the PCC seed, the dorsal attentional regions only occured in the case of the spatially distributed regions (See Fig. 3, X=38).

Overall, these findings strongly support the existence of multiple states either in the case of the spatially distributed parcels or the case of spatially contiguous regions.

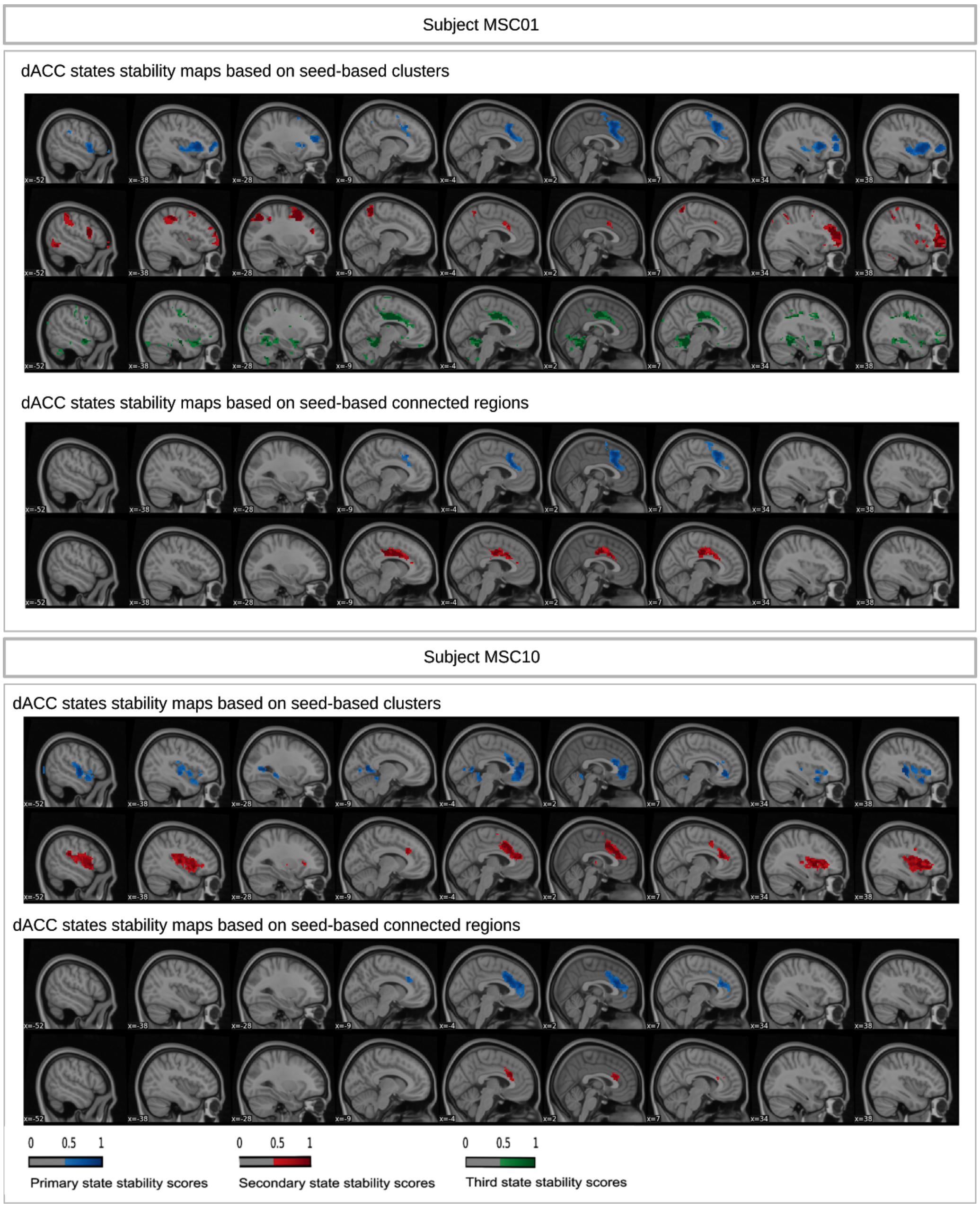


**Figure 2. The spatially contiguous regions were overlapping with the spatially distributed parcels surrounding the seed in the state stability maps of the dACC seed**. The minimum size of regions in the spatially contiguous regions was fixed to 50 voxels. The number of replications of seed-based parcellations = 30. The cluster size threshold = 5%. The seed-based parcellations were clustered into 12 clusters. The smoothing kernel = 6 mm, the cluster size threshold = 10%, the Dice threshold = 0.3, the number of timepoints in the window length = 100. Subject MSC01 and Subject MSC10 of the Midnight scan club dataset were included.

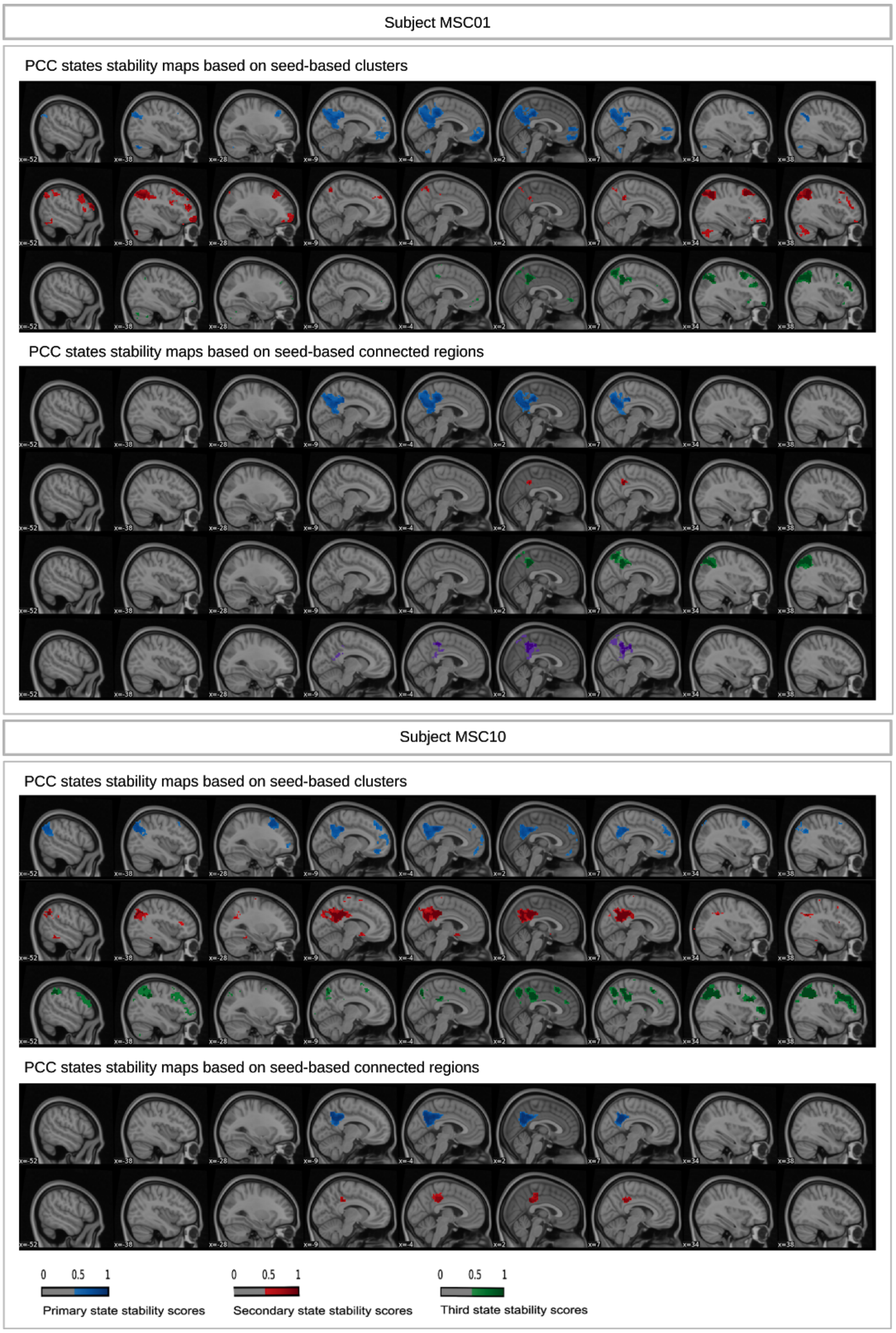


**Figure 3.** The spatially contiguous regions were overlapping with the spatially distributed parcels surrounding the seed in the state stability maps of the PCC seed. The minimum size of regions in the spatially contiguous regions was fixed to 50 voxels. The number of replications of the seed-based parcellations = 30. The cluster size threshold = 5%. The seed-based parcellations were clustered into 12 and 50 clusters. The smoothing kernel = 6

mm, the cluster size threshold = 10%, the Dice threshold = 0.3, the number of timepoints in the window length = 100. Subject MSC01 and Subject MSC10 of the Midnight scan club dataset were included.

#### Supplement 3

##### Spatial similarity of the seed-based parcellations

We aimed to investigate the spatial similarity of the seed-based parcellations used to generate the dynamic state stability maps. To this end, we reported the Dice scores between pairs of seed-based parcellations across seeds and subjects. Three seed voxels were investigated including PM-VIS, dACC and PCC. Our results showed that some seed-based parcellations were very highly correlated with near perfect Dice scores (~=1) and that Dice scores were around the median ~ 0.23 for the PM-VIS and the PCC seeds while the dACC had a lower median with Dice score = 0.2 (See Fig. 4).

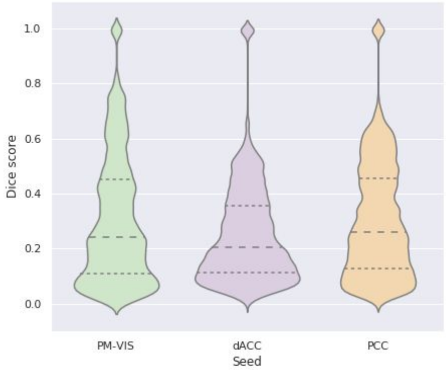


**Figure 4. Spatial similarity between seed-based parcellations in terms of Dice scores.** We included the Dice scores associated with seed-based parcellations. We studied three seed voxels from the PM-VIS, dACC and PCC subnetworks. The smoothing kernel = 6 mm. Number of timepoints in the window length = 100. Ten subjects from the Midnight scan club dataset were included.

#### Supplement 4

##### 4.1 Reproducibility of our dynamic states of parcellations was insensitive to resolution, yet some spatial states were consistent for some subjects and variable for others

We aimed to investigate the effect of the resolution; i.e. the number of clusters, in the seed-based parcellations on the reproducibility of dynamic states of parcellations. To this end, we computed the within- and the between-subject reproducibility. Our results showed that the within-subject reproducibility of the dynamic states of parcellations was higher than the between-subject reproducibility both in the cases of 12 and 50 clusters (See Fig. 5). For instance, the PCC median of the within-subject reproducibility was about 0.9 in both 12 and 50 cluster cases. Moreover, the between-subject reproducibility distributions were highly overlapping across the three seeds.

We also observed that the 50 cluster-based states had overlapping regions with the 12 cluster-based states at the level of voxels surrounding the seed for some subjects and states. For instance, subject MSC01 and the dACC primary states overlapped for the dACC regions surrounding the seed with a larger spatial distribution in the case of the 12 clusters (Fig. 6, X=2). Similarly, the secondary dACC states and subject MSC03 had some overlap in the dACC regions surrounding the seed even though the 12 clusters regions were more lateralized toward the motor regions (See Fig. 7, X=2). Moreover, we observed that the 50 cluster-based states had an overlap with the 12 cluster-based states at the level of the distributed regions throughout the brain. For instance, the dACC primary states and subject MSC01 had an overlap at the level of the insular regions (See Fig. 6, X=38). Similarly, both the dACC secondary states of subject MSC06 involved the insular regions (See Fig. 7, X=38).

Conversely, some subjects and states had either a negligible overlap, inconsistencies in the regions or absence of the 12 clusters based-states. For instance, the 50-clusters primary state in the case of the dACC and subject MSC07 had a very small dACC region compared to the 12-clusters primary state (see Fig. 6, X=4). Also, the 12-clusters secondary state and subject MSC08 had lateralized dACC regions towards the motor regions while the 50-clusters secondary state involved regions from the visual cortex (See Fig. 7, X=4). Also, most states of the third states were inconsistent across scales (See Fig. 8).

Overall, the reproducibility of our dynamic states of parcellations was insensitive to the scale, yet spatial states were consistent for some states and subjects and variable across others.

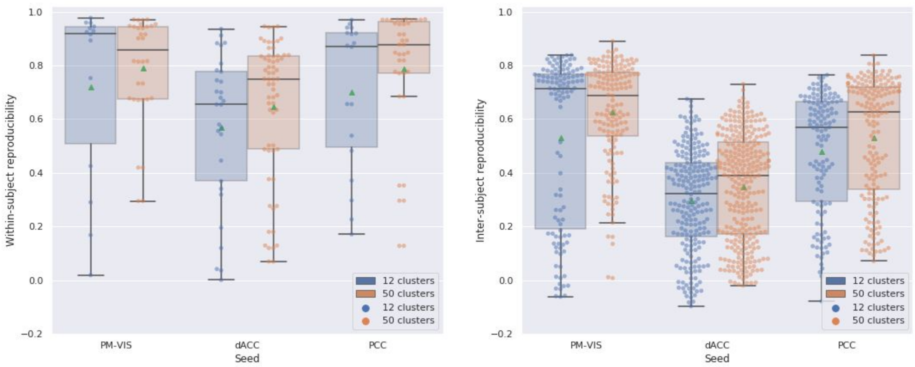


**Figure 5**. **Within-subject reproducibility of dynamic states was higher at the within-subject level than the between-subject level across scales.** We studied three seed voxels: PM-VIS, dACC and PCC. The number of replications of our Dypac algorithm replications with different seeds = 30. Number of replications of seed-based parcellations =30. The cluster size threshold = 5%. The seed-based parcellations were clustered into 12 and 50 clusters. The smoothing kernel = 6 mm, the Dice threshold = 0.3. The number of timepoints in the window length = 100. Ten subjects of the Midnight scan club dataset were included.

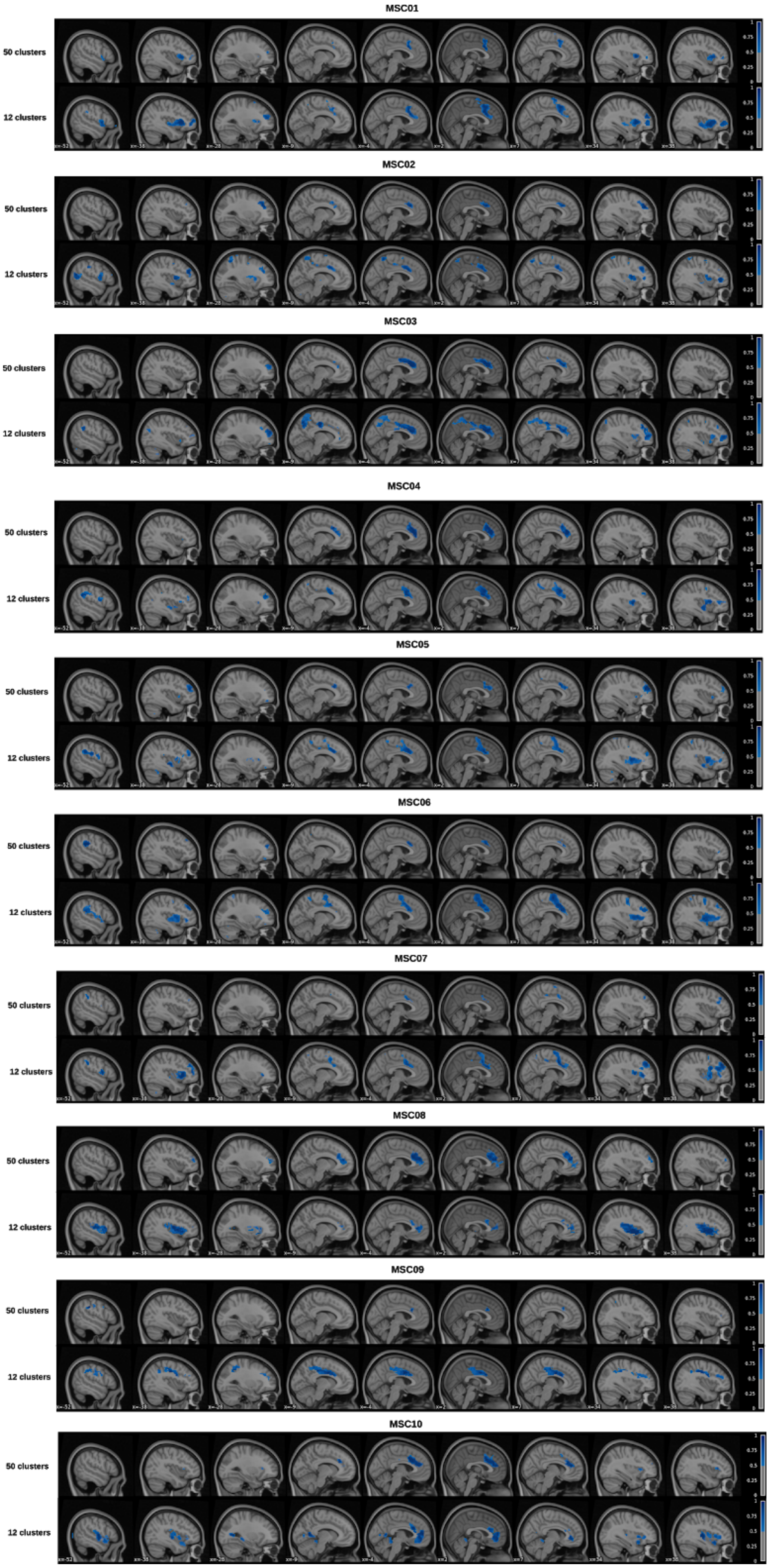
MSC01
50 clusters
12 clusters
MSC02
50 clusters
12 clusters
MSC03
50 clusters
12 clusters
MSC04
50 clusters
12 clusters
MSC05
50 clusters
12 clusters
MSC06
50 clusters
12 clusters
MSC07
50 clusters
12 clusters
MSC08
50 clusters
12 clusters
MSC09
50 clusters
12 clusters
MSC10
50 clusters
12 clusters

**Figure 6. Primary states were overlapping across scales for some subjects in the case of the dACC seed**. We studied three seed voxels: PM-VIS, dACC and PCC. The number of replications of our Dypac algorithm replications with different seeds = 30. The number of replications of the seed-based parcellations for each replication = 30. The cluster size threshold = 5%. The seed-based parcellations were clustered into 12 and 50 clusters. The smoothing kernel = 6 mm, the Dice threshold = 0.3. The number of timepoints in the window length = 100. Ten subjects of the Midnight scan club dataset were included.

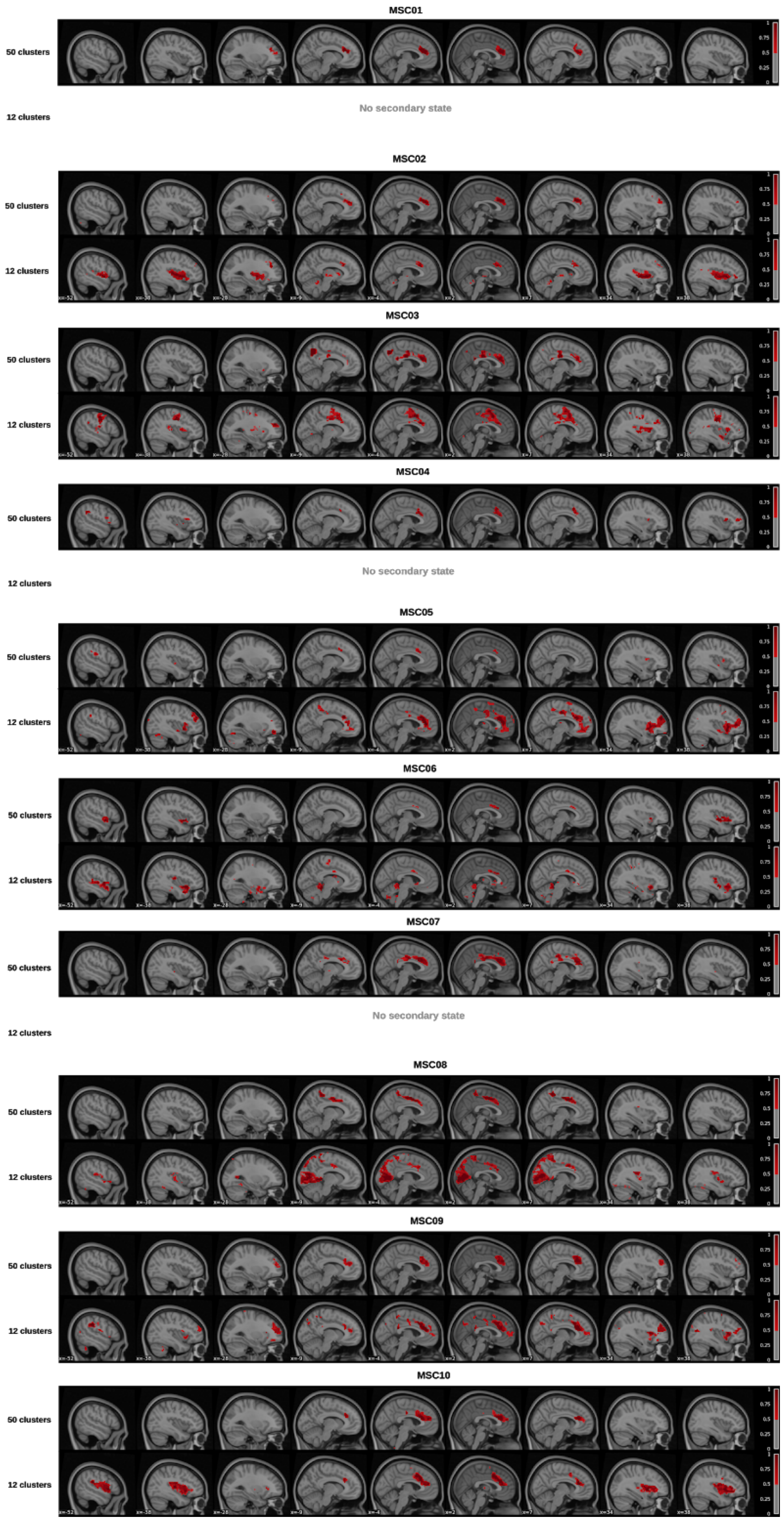


**Figure 7. Secondary states were overlapping across scales for some subjects and states and variable across others in the case of the dACC seed**. We studied three seed voxels: PM-VIS, dACC and PCC. The number of replications of the seed-based parcellations

for each replication = 30. The cluster size threshold = 5%. The seed-based parcellations were clustered into 12 and 50 clusters. The smoothing kernel = 6 mm, the Dice threshold = 0.3. The number of timepoints in the window length = 100. Ten subjects of the Midnight scan club dataset were included.

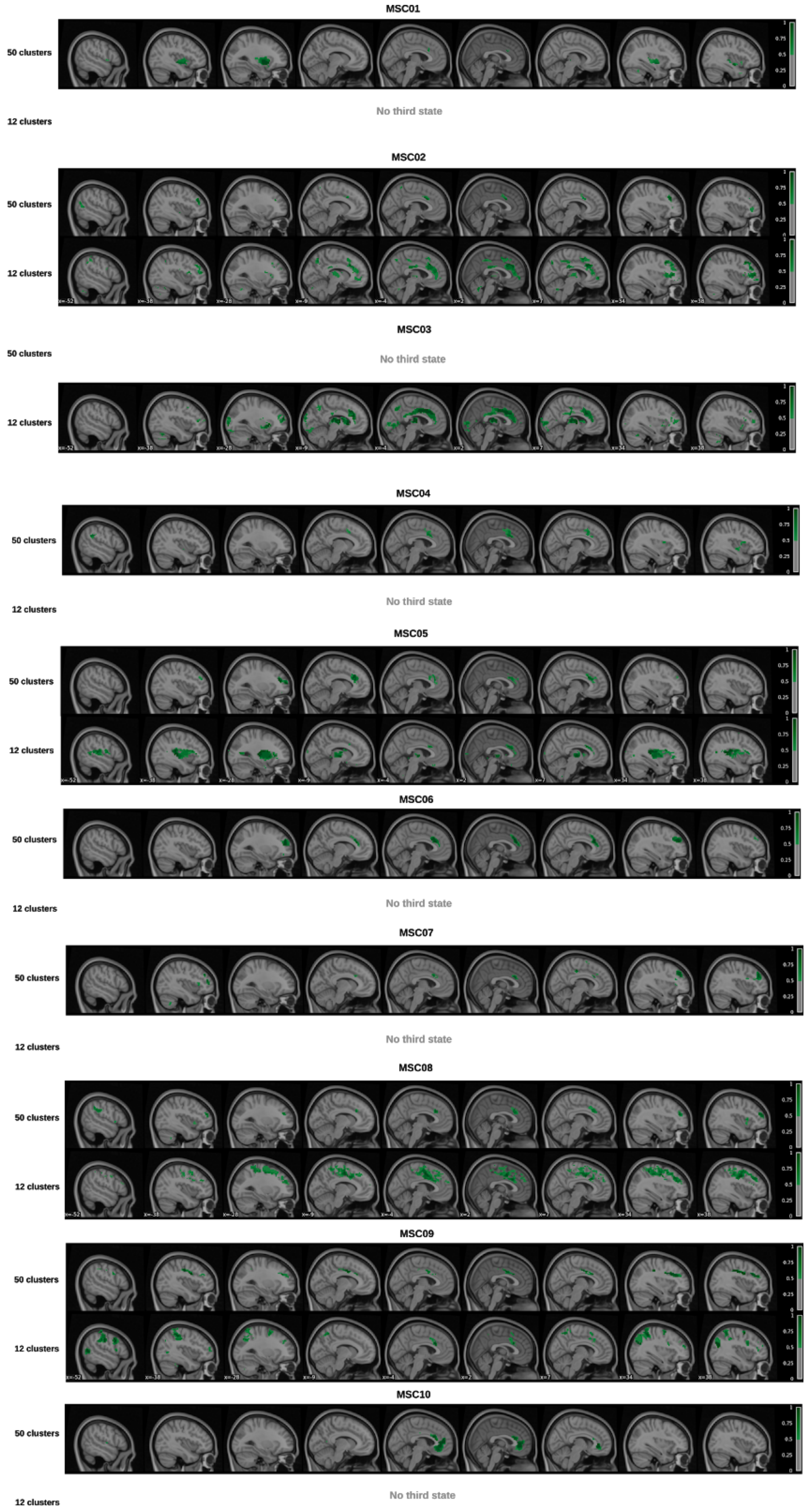


**Figure 8. Third dACC dynamic states were inconsistent across scales**. We studied three seed voxels: PM-VIS, dACC and PCC. The number of replications of our Dypac

algorithm replications with different seeds = 30. The number of replications of the seed-based parcellations for each replication = 30. The cluster size threshold = 5%. The seed-based parcellations were clustered into 12 and 50 clusters. The smoothing kernel = 6 mm, the Dice threshold = 0.3. The number of timepoints in the window length = 100 timepoints. Ten subjects of the Midnight scan club dataset were included.

##### 4.2 Dynamic states can be identified at different time scales

We aimed to investigate the effect of the window length on the reproducibility of the dynamic states of parcellations. Thus, we computed the within- and the between-subject reproducibility for different window lengths; i.e. the number of timepoints = {30, 50, 100, 200} in window lengths for the ten subjects of the Midnight scan club dataset. Our results showed that within-subject reproducibility was higher than between-subjects reproducibility for all window length values (see Fig. 9). For instance, in the case of 50 timepoints, within-subject reproducibility scores had a Pearson correlation median > 0.83 across seeds. However, between-subjects reproducibility scores had a correlation median < 0.78 across seeds (See Fig. 9). Similarly, in the case of 200 timepoints, within-subject reproducibility scores had a Pearson correlation score > 0.7, however, between-subjects reproducibility scores had a median correlation score < 0.7 (See Fig. 9).

For different number of timepoints in the sliding windows, the studied subnetworks were multistate in the case of the dACC and the PCC seeds while the PM-VIS was monostate for most subjects and at multiple time scales. However, we observed the absence of states for some subjects in the case of 30 timepoints. The sensitivity of the k-Means algorithm to local minima due to few time samples may be at the origin of this limitation.

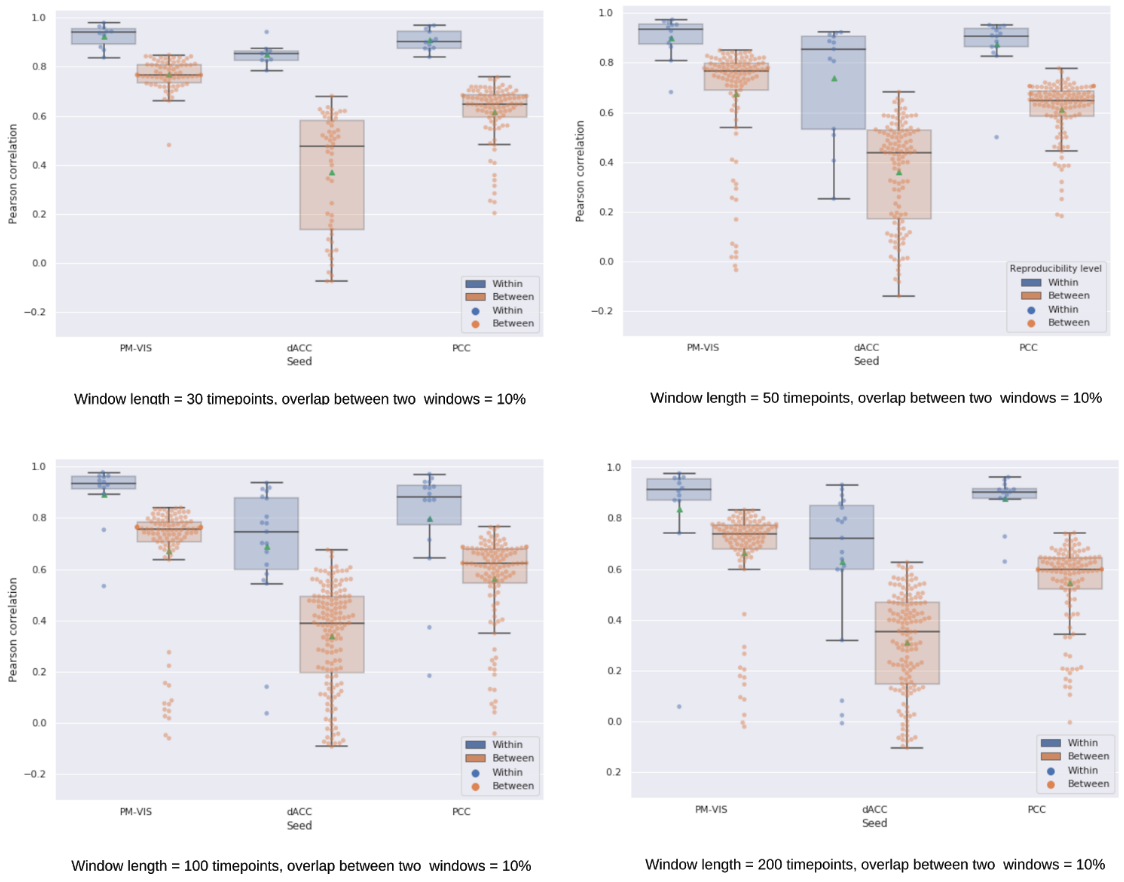


**Figure 9. Within-subject reproducibility scores were higher than between-subject reproducibility scores for most dynamic states of parcellations.** In the case of 30 timepoints, some subjects did not capture any state. We studied three seed voxels: PM-VIS, dACC and PCC. The number of replications of seed-based parcellations = 30. The seed-based parcellations were clustered into 12 clusters. The smoothing kernel = 6 mm, the cluster size threshold = 10%, the Dice threshold = 0.3. Ten subjects of the Midnight scan club dataset were included.

##### 4.3 Subnetworks were multistate with a dominant primary state for different window lengths

We investigated the distribution of the states dwell time for different number of timepoints per window; i.e., number of timepoints = {30, 50, 100, 200}. Three seeds from the PM-VIS, the dACC and the PCC subnetworks were investigated. Our results showed that the dACC and the PCC seeds were multistate for different number of timepoints in a single window. For instance, the dwell time of the dACC seed had a maximum of three states in the case of 30 time points, three states in the case of 50 timepoints, five states in the case of 100 time points and four states in the case of 200 time points. Moreover, the primary state was always a dominant state with an important difference between its dwell time and the dwell time of the secondary state. For instance, in the case of 50 time points and the dACC seed, the primary state had a dominant dwell time with a median = 38% over a median dwell time equal to 11% in the case of the secondary state. Similarly, the dwell time of the PCC seed had a maximum of three, four, four and three states, respectively for the 30, 50, 100 and 200 timepoints. The dominant primary state in the case of 50 timepoints and the PCC seed had a median dwell time = 42% over a median dwell time = 11% in the case of its secondary state. Unlike the dACC and the PCC, the PM-VIS seed was monostate for most subjects and up to three states for few subjects. Its secondary and third states had very low dwell time (~10%) compared to the primary state (See Fig. 10). Altogether, these results suggested that the dynamic states of parcellations were multistate for the dACC and the PCC seeds and mono-state for the PM-VIS seed for most subjects and multistate for few subjects, regardless of the window time length.

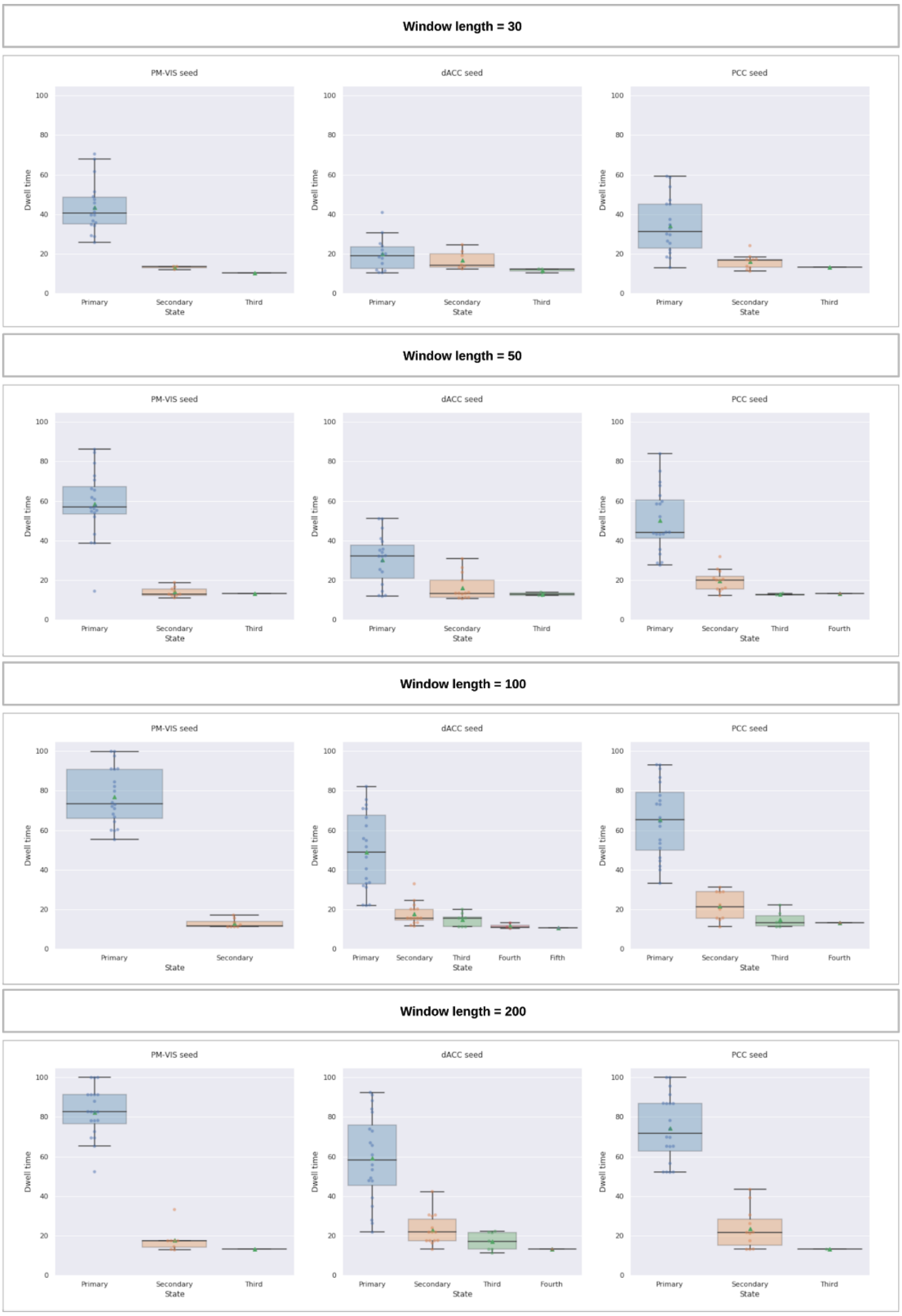


**Figure 10. The number of dynamic states and their dwell time was insensitive to different window lengths**. We reported the dwell time from both sets of independent sessions. We studied three seed voxels including the PM-VIS, the dACC and the PCC. The number of replications of seed-based parcellations = 30. The seed-based parcellations were clustered into 12 clusters. The smoothing kernel = 6 mm, the cluster size threshold = 10%, the Dice threshold = 0.3. Ten subjects of the Midnight scan club dataset were included.

##### 4.4 Dynamic states reproducibility was insensitive to low cluster size thresholds

We aimed to investigate the impact of different cluster size thresholds on the reproducibility of dynamic states of parcellations. Thus, we reported the within- and between-subjects reproducibility for different cluster size thresholds as the percentage of the seed-based parcellations included in a given state over their total number. The results of the lowest cluster size threshold; i.e. score = 5%, showed large distribution of reproducibility scores both at the within- and the between-subjects levels compared to higher cluster size thresholds; i.e. score > 10%. Still, the median of the within-subject reproducibility was higher than the between-subject reproducibility for all cluster size thresholds and all seeds (See Fig. 11). These results demonstrated the insensitivity of the reproducibility of the dynamic states of parcellations to the cluster size threshold.

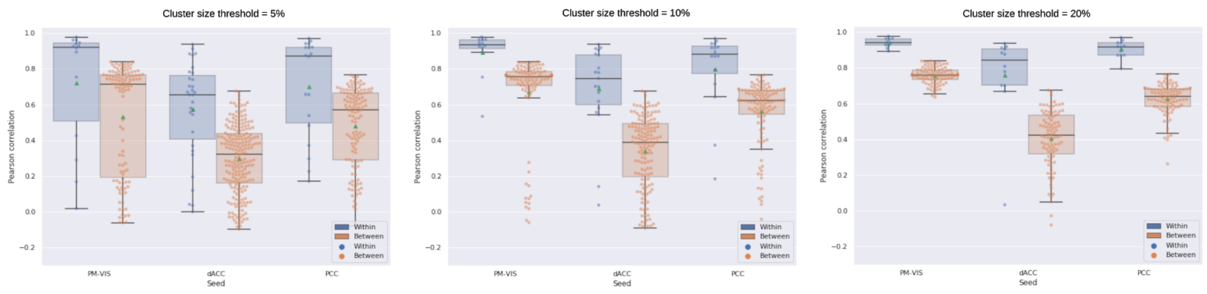


**Figure 11. Dynamic states reproducibility was insensitive to cluster size threshold**. We studied three seed voxels including the PM-VIS, the dACC and the PCC. The number of replications of seed-based parcellations = 30. The seed-based parcellations were clustered into 12 clusters, the smoothing kernel = 6 mm, the Dice score threshold = 0.3. The number of timepoints per window = 100. Ten subjects of the Midnight scan club dataset were included.

##### 4.5 The PCC and the dACC subnetworks were multistate and the PM-VIS was monostate for most subjects across different cluster size thresholds

We investigated the distribution of the states dwell time for different values of cluster size thresholds = {5%, 10%, 20%}. Three seeds from the PM-VIS, the dACC and the PCC subnetworks were investigated. Our results showed that the dACC and the PCC seeds were multistate across cluster size thresholds. For instance, the dwell time of the dACC seed had a maximum of five states in some subjects when the cluster size thresholds were equal to 5% and 10% while it had only up to three states in the case of 20% cluster size threshold. Moreover, the primary state was always a dominant state with an important difference between its dwell time and the dwell time of the secondary state of all the values of cluster size thresholds. For instance, in the case of a cluster size threshold equal to 5%, the dACC seed and the primary state had a dominant dwell time with a median = 43% over a median dwell time = 13% in the case of the secondary state (See Fig. 12). Altogether, these results suggested that the dynamic states of parcellations were multistate for the dACC and the PCC seeds. Dynamic states of parcellations were mono-state in the case of the PM-VIS seed for most subjects and multistate for few subjects, regardless of the cluster size threshold.

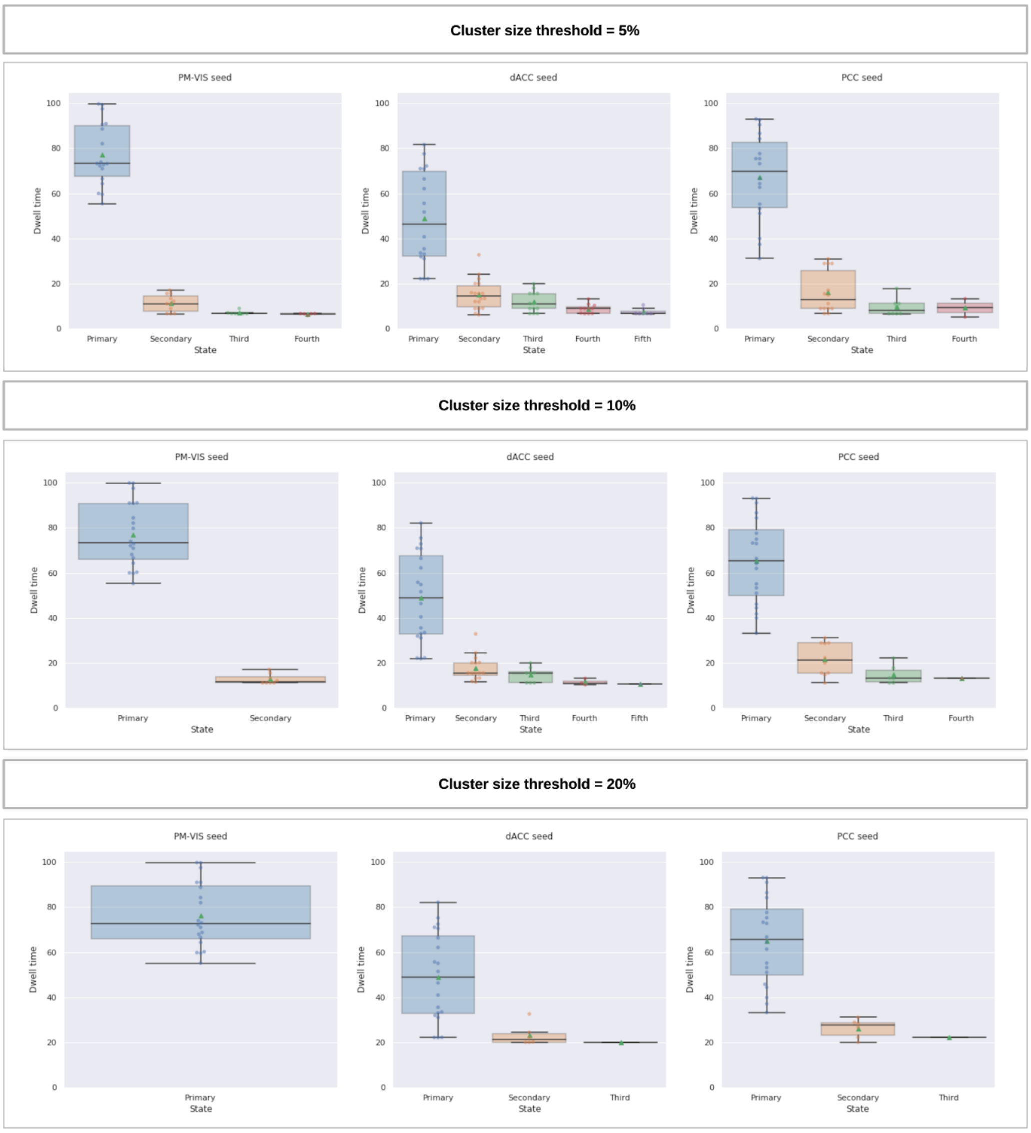


**Figure 12. The number of dynamic states and their dwell time was insensitive to different cluster size thresholds**. We studied three seed voxels including the PM-VIS, the dACC and the PCC. The number of replications of the seed-based parcellations = 30. The seed-based parcellations were clustered into 12 clusters. The smoothing kernel size = 6 mm, the Dice score threshold = 0.3. The number of timepoints per window = 100. Ten subjects of the Midnight scan club dataset were included.

#### 4.6 Reproducibility of dynamic states of parcellations in the case of different smoothing kernels

We aimed to investigate the impact of the spatial smoothing on the spatial state maps. Thus, we generated the dynamic states of parcellations for different smoothing kernels; i.e., 4mm, 6mm and 8mm, and we reported the within- and between-subjects reproducibility. Our results showed that all within-subjects reproducibility were higher than between-subjects reproducibility. For instance, the median of within-subject reproducibility scores were higher than 0.8 Pearson correlation for all seeds across different smoothing kernels while the between-subjects reproducibility were lower than 0.8 Pearson correlation for all seeds and all smoothing kernels (See Fig. 13). These findings suggested the insensitivity of our dynamic states of parcellations to the kernel size.

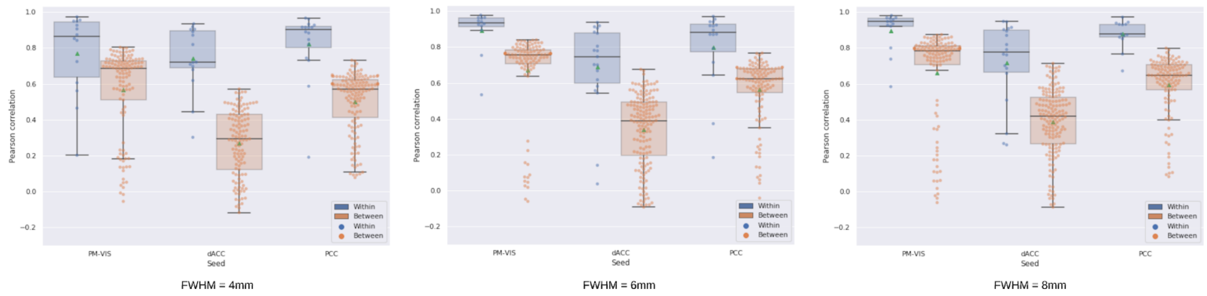


**Figure 13. The within- and between-subjects reproducibility of our dynamic states of parcellations were insensitive to the smoothing kernel size**. We studied three seed voxels: PM-VIS, dACC and PCC. The number of replications of seed-based parcellations for each replication=30. The seed-based parcellations were clustered into 12 clusters, the cluster size threshold = 10%, the Dice threshold = 0.3. The number of timepoints per window = 100. Ten subjects of the Midnight scan club dataset were included.

#### 4.7 The PCC and the dACC subnetworks were multistate for different smoothing kernels and the PM-VIS was mono-state for most subjects

We aimed to get a better understanding of the dwell time distribution of the dynamic states of parcellations for different smoothing kernels. Our results showed the existence of many states across the three studied subnetworks; i.e. dACC, PCC and PM-VIS seeds. For instance, the PM-VIS had three, two and three states, respectively for 4mm, 6mm and 8mm smoothing kernels. However, the dwell times of the secondary and the third states of the PM-VIS subnetwork occured for a few subjects and their dwell time was very low; i.e. dwell time ~10%. That suggested the PM-VIS was mono-state for most subjects and multistate for some subjects with a dominant primary state. Conversely, the dACC and the PCC secondary states had much lower dwell time than the primary state; i.e. dwell time > 10%, for different smoothing kernels. These findings suggested the PCC and the dACC subnetworks were multistate while the PM-VIS was mono-state for different smoothing kernels.

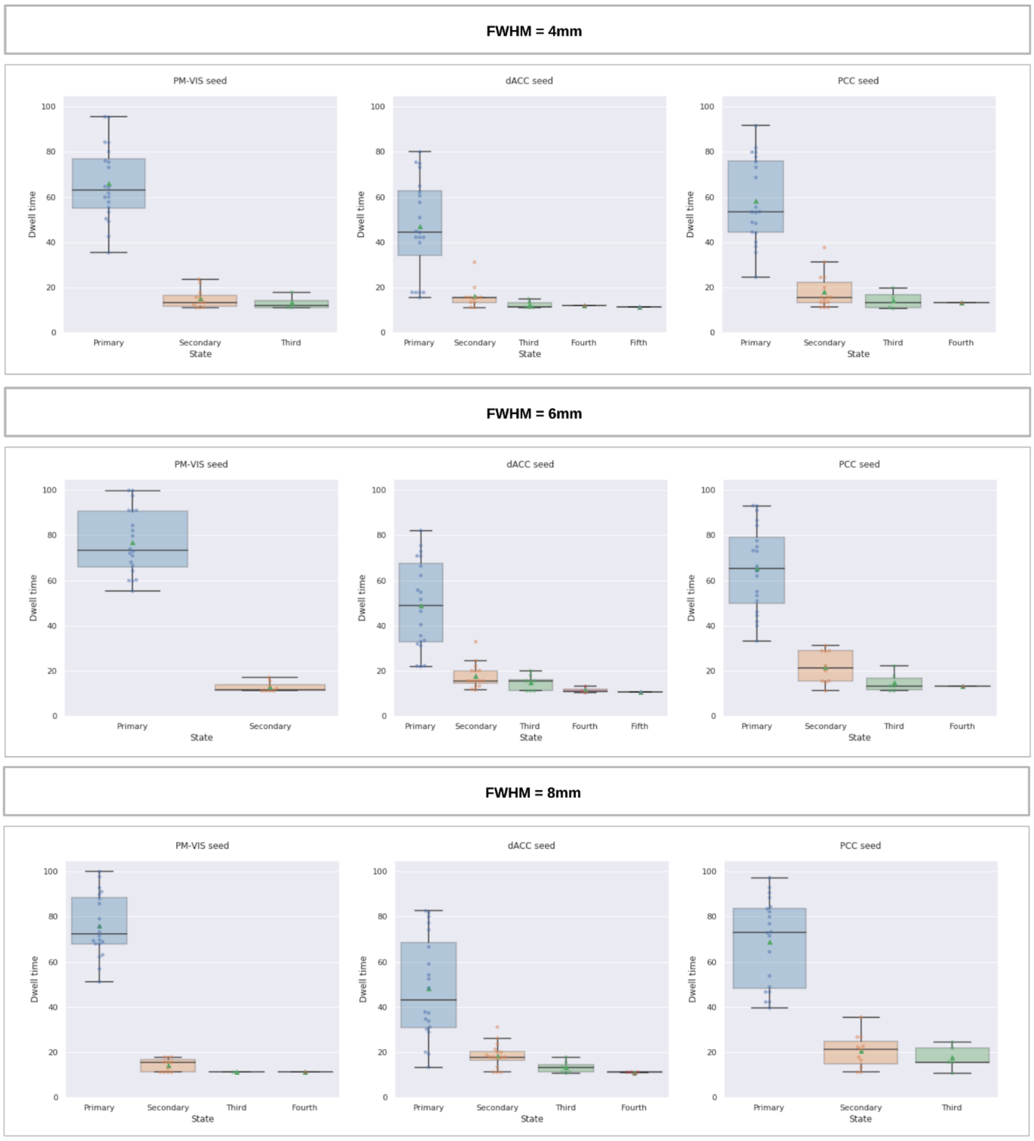


**Figure 14. The number of dynamic states and their dwell time was insensitive to the smoothing kernel**. We studied three seed voxels including the PM-VIS, the dACC and the PCC. The number of replications of seed-based parcellations = 30. The cluster size threshold = 10%. The seed-based parcellations were clustered into 12 clusters, the Dice threshold = 0.3, the number of timepoints per window = 100. Ten subjects of the Midnight scan club dataset were included.

##### 4.8 The reproducibility of the dynamic states of parcellations was not sensitive to changes in the seeds coordinates

We aimed to investigate the reproducibility of many seed voxels from the visual, the dACC and the PCC subnetworks to verify the generalizability of the conclusions associated with the studied subnetworks. We manually picked 15 different seeds from each subnetwork including boundaries (See MNI coordinates of the chosen seeds in table 1). We computed their within-subject and between-subjects reproducibility scores. Our results showed that within-subject reproducibility outperformed the between-subjects reproducibility scores in terms of the Pearson correlation. For instance, the within-subjects reproducibility scores had a median correlation > 0.9 in the case of the PCC seed while all between-subjects reproducibility scores had a median correlation < 0.79 (See Fig. 15). These results suggested that within-subjects reproducibility of our dynamic states of parcellations was higher than between-subjects reproducibility for all the studied seeds from the PM-VIS, the dACC and the PCC subnetworks.

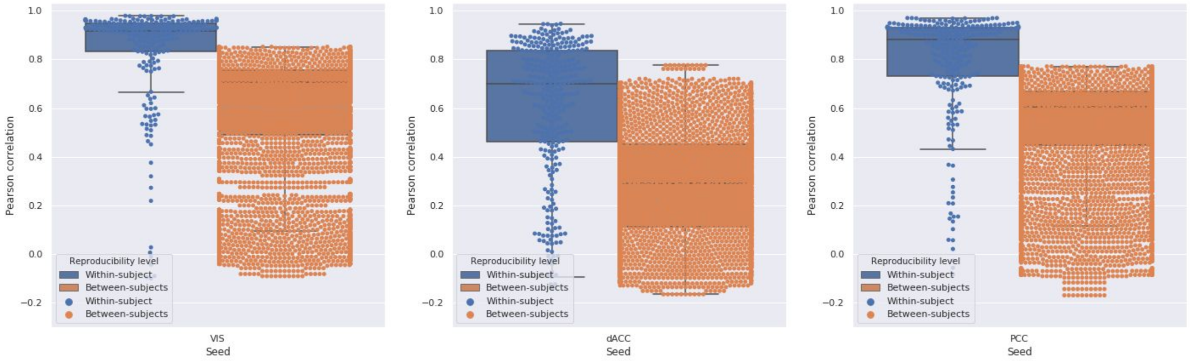


**Figure 15. Within- and between-subjects reproducibility of our dynamic states of parcellations for 15 seeds per subnetwork**. We studied three seed voxels including the PM-VIS, the dACC and the PCC. The number of replications of the seed-based parcellations was equal to five. The seed-based parcellations were clustered into 12 clusters. The smoothing kernel size = 6 mm, the cluster size threshold = 10%, the Dice threshold = 0.3, the number of timepoints per window = 100. Ten subjects of the Midnight scan club dataset were included.

##### 4.9 Within-subjects reproducibility was insensitive to the number of replications of the seed-based parcellations

To evaluate the impact of the number of replications of the seed-based parcellations, we computed the within-subjects reproducibility of the dynamic states of parcellations in the case of 1, 5 and 30 seed-based parcellations. Different initializations of the random number generator were used for different k-Means parcellations. Our results showed very similar distributions across replications. For instance, the within-subjects reproducibility medians were all aligned to very close scores for all seeds (See Fig. 16). These findings suggested that the variability across seeds in the k-Means parcellations were negligible in the context of our Dypac algorithm.

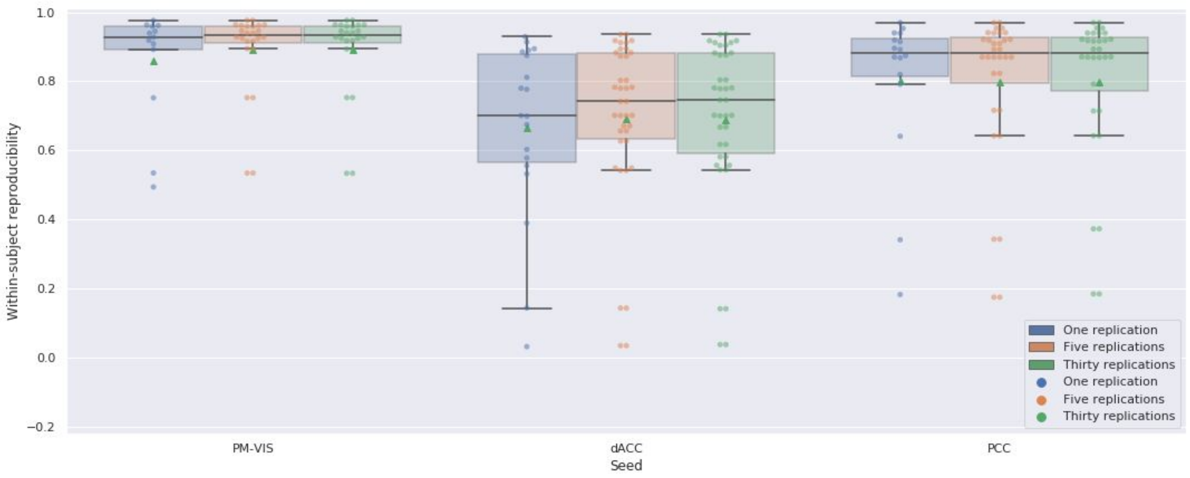


**Figure 16. The within-subject reproducibility of our dynamic states of parcellations was insensitive to the number of replications of the seed-based k-Means parcellations**. We studied three seed voxels including the PM-VIS, the dACC and the PCC. The seed-based parcellations were clustered into 12 clusters. The smoothing kernel = 6 mm, the cluster size threshold = 10%, the Dice threshold = 0.3, the number of timepoints per window = 100. Ten subjects of the Midnight scan club dataset were included.

### Supplement 5

#### Dynamic states suppressed differences within-sessions versus across days in the case of resting state functional MRI data

We aimed to investigate the within-sessions versus the between sessions effect across days on the identified dynamic states. To this end, we computed the probability of a given state to be associated with two sliding windows either from the same session or from different sessions (across days). Our results showed highly overlapping probability distributions for the sliding windows of the same state to fall into the same session or different sessions (See Fig. 17). Thus, there was no substantial impact of the differences in the brain activity across sessions on the identified dynamic states.

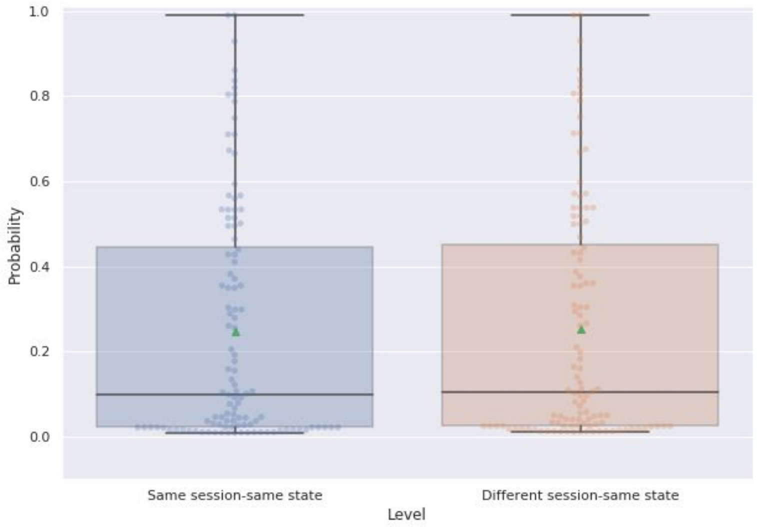


**Figure 17. Dynamic states occurred with the same probability either across days or across sessions.** For each state, we computed the probability of two sliding windows to be in the same session or in different sessions. We studied three seed voxels including the PM-VIS, the dACC and the PCC. The number of seed-based parcellations = 5. Ten subjects of the Midnight scan club dataset were included.

#### Supplement 6

##### Different dynamic states had different temporal dynamics

We aimed to quantify the synchrony between the dynamics of the different states of parcellations, from different seeds and over time. To this end, we computed the Adjusted Rand Index (ARI) similarity scores between one-hot encoding of states associated with seed-based parcellations (over sliding windows) as a measure of the synchrony between spatial states over time. The sliding windows were ordered chronologically. The seed-based one-hot parcellation dynamics was the binary representation in which 1 value indicated the sliding window was included in the state and 0 otherwise, for a given sliding window. We computed the ARI between one-hot temporal dynamics associated with different seeds for five sets of sessions (within-set) and between two sets of five independent sessions (between-sets). This allowed us to quantify to which extent the spatial spatial patterns (states) associated with different seed subnetworks were involved, at the same time (over sliding windows) both within- and between-sets. No substantial association between sets was expected, as these sessions were acquired independently making inter-set synchronization unlikely, with the possible exception of habituation, fatigue or stress effects accruing systematically at the beginning or the end of a session. The low ARI scores within- and between-sets showed there was no synchrony between the states of different pairs of seeds both at the within-set and between-sets. That is, the states associated with one seed did not occur at the same time as the states of another seed. Also, we observed that the within-set scores were slightly higher than between-sets scores in the case of the PCC/PM-VIS pair. For instance, for the three pairs of seeds (i.e. dACC/PCC, dACC/PM-VIS,

PCC-PM-VIS), the average ARI score ~0.02. However, The PCC/PM-VIS pair had an average ARI of 0.05 (See Fig. 18). These findings demonstrated the existence of different temporal dynamics for the states associated with different seed subnetworks.

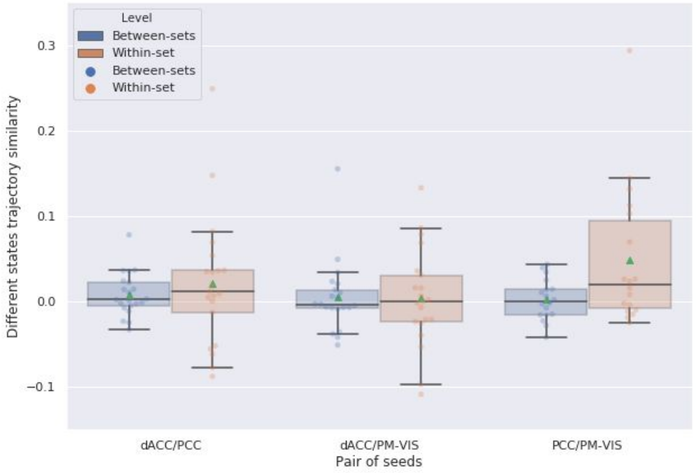


**Figure 18**. **Dynamic states of parcellations had different temporal dynamics**. The ARI similarity scores were computed between the seed-based parcellations associated with different pairs of seed subnetworks. Each seed-based parcellation was a chronologically ordered sliding window parcellation from the MSC sessions. We computed the ARI either between the states from at the within-set level or at the between-sets level. We maximized the ARI scores between two distinct states associated with two different seed subnetworks (i.e. dACC/PCC, dACC/PM-VIS, PCC/PM-VIS). We replicated the Dypac algorithm 30 times. The number of seed-based parcellations = 5. The seed-based parcellations were clustered into 12 clusters. The smoothing kernel = 6 mm, the cluster size threshold = 10%, the Dice threshold = 0.3, the number of timepoints per window = 100. Ten subjects of the Midnight scan club dataset were included.

| **Visal sbnetwork seeds** | **dACC subnetwork seeds** | **PCC subnetwork seeds** |
|---|---|---|
| (-6.0, -79.0, -8.0) | (-3.0, 14.0, 34.0) | (0.0, -43.0, 34.0) |
| (0.0, -76.0, 10.0) | (-3.0, 14.0, 40.0) | (-3.0, -46.0, 37.0) |
| (0.0, -79.0, 19.0) | (-3.0, 26.0, 34.0) | (0.0, -46.0, 43.0) |
| (0.0, -70.0, 19.0) | (-3.0, 29.0, 31.0) | (6.0, -46.0, 34.0) |
| (-6.0, -91.0, 19.0) | (-12.0, 32.0, 22.0) | (0.0, -37.0, 40.0) |
| (15.0, -82.0, 7.0) | (-9.0, 23.0, 22.0) | (-3.0, -37.0, 34.0) |
| (15.0, -94.0, 1.0) | (-6.0, 11.0, 37.0) | (-3.0, -49.0, 34.0) |

| | | |
|---|---|---|
| (15.0, -64.0, 7.0) | (-6.0, 35.0, 25.0) | (6.0, -52.0, 13.0) |
| (15.0, -76.0, 31.0) | (-6.0, 38.0, 16.0) | (6.0, -34.0, 34.0) |
| (15.0, -64.0, 13.0) | (-6.0, 17.0, 28.0) | (6.0, -64.0, 34.0) |
| (-18.0, -88.0, 22.0) | (-9.0, 8.0, 34.0) | (6.0, -46.0, 28.0) |
| (-18.0, -79.0, -8.0) | (-9.0, 26.0, 25.0) | (-3.0, -49.0, 37.0) |
| (-18.0, -64.0, -5.0) | (-3.0, 26.0, 19.0) | (-3.0, -52.0, 19.0) |
| (-18.0, -64.0, 16.0) | (6.0, 26.0, 22.0) | (-9.0, -40.0, 34.0), |
| (-18.0, -85.0, 34.0) | (6.0, 8.0, 28.0) | (12.0, -52.0, 28.0) |

**Table 1. Extended list of MNI seed coordinates from the three subnetworks: Visual subnetwork, dACC subnetwork and PCC subnetwork.**
